# A network-based framework for detecting communities of similar neural spike trains

**DOI:** 10.64898/2026.09.03.749154

**Authors:** Indranil Ghosh, Áine Byrne

## Abstract

Recent advances in large-scale electrophysiological recording technologies now allow simultaneous measurement of spike trains from ensembles of neurons. A central challenge in mathematical neuroscience is therefore to identify functional assemblies and their collective organisation directly from this data. The goal of this work is to devise a method to infer the collective organisation of the neurons from their spiking activity. We construct weighted functional networks from neural spike trains using the van Rossum distance to quantify pairwise similarity. The functional network captures similarities in neuron firing patterns and provides a representation of their collective organisation. We hypothesise that similar neuronal assemblies will appear as clustered communities in the network, and employ the Louvain algorithm to detect such assemblies. We validate our approach using synthetic spike train data and simulated data from a stochastic block model of Leaky Integrate and Fire neurons subjected to external Poisson drives, where the ground truth is known. Finally, we apply our approach to large-scale recordings from the Allen Institute Visual Coding: Neuropixels dataset. We find that our methodology works well as long as a sufficient amount of data is available and the temporal structure is strong relative to the noise.

## 1 Introduction

In modern neuroscience research, one of the central goals is to understand how the collective activity of large assemblies of neurons leads to the emergence of cognition and behaviour. Groups of neurons typically organise into coordinated assemblies rather than acting as independent single neuron responses [1]. They are temporal objects in general, meaning that the neurons from a functional assembly are expected to display similar spike timing patterns even though their exact spike timings are not exactly matching. For example, hippocampal “place cells” form temporally coordinated ensembles whose activity can be directly related to encoding an animal’s reference. The reactivation of these temporal sequences during sleep and resting wakefulness has been related to mechanism for consolidating spatial memories [2]. Furthermore, coordinated firing among neuron populations has been linked to encoding and transmission of sensory information [3]. In the motor cortex, assemblies of neurons exhibit coordinated collective dynamics during reaching movements [4], suggesting that motor behaviour is encoded collectively across neural populations rather than by individual neurons alone. Similar neural-population level structured activity is also observed during resting state dynamics. For example, the default mode network exhibits coherent spontaneous activity across distributed brain regions even in the absence of external tasks [5]. Also, spontaneous activity in the visual cortex are organised according to visual processing streams [6]. All of this evidence suggests that neural assemblies may point towards an intrinsic network organisation and latent functional architecture. Understanding how coordinated neural activity gives rise to functional organisation has become a central goal of modern network neuroscience research [7]. Identifying and characterising these assemblies according to their functional properties from large scale neural recording datasets is an important challenge for researchers studying neural data.

Despite the availability of large amounts of spike-train data from electrophysiological recordings, identifying a functional group of neurons according to their activity patterns is still a difficult task. Existing approaches have addressed this by reconstructing functional or effective connectivity networks from neural recordings [8] and subsequently applying network-theoretic analyses to identify functional assemblies [9, 10]. More recent methods have focused on structured representations of population spike patterns and their geometric relationships [11]. Many methods rely on correlation-based [12, 13, 14, 15] or statistical methods [16, 9] that do not fully exploit precise spike timing information. Also, validation against known ground-truth assemblies is often limited, making it difficult to determine the robustness of proposed methodologies. Furthermore, not much attention is given to assessing functional community detection as function of biologically relevant parameters like the inter-spike intervals (ISI), spike-time jitter, recording duration, network size, and the temporal scale at which spike trains are compared. In our work we address these challenges by developing a framework called the **vR-FCD** (*van Rossum-Functional Community Detection*) that combines the *van Rossum* [17, 18, 19] similarity measure with network based community detection (Louvain method [20]) to find functional assemblies of neurons. Unlike previous studies relying on thresholding to construct binary networks [9], our method preserves pairwise similarity weights by constructing a fully connected weighted functional network. Application of the multilevel Louvain method is a step forward compared to applying earlier modularity optimisation based community detection like Girvan and Newman [21].

The remainder of the paper is organised as follows: In § 2 we describe the vR-FCD framework detailing its steps. Then in § 3 we develop a synthetic spike-activity dataset with controllable planted community structures and apply the framework. In § 4 we extend our application of the vR-FCD frameowork to a stochastic block model (SBM) network of conductance based Leaky Integrate and Fire (LIF) neurons subjected to external Poisson inputs. In § 5, we finally apply the framework to the Neuropixels recordings from awake wild-type mice provided by Allen Brain Institute and perform robustness tests of our method besides studying the correspondence between communities detected using the vR-FCD framework and the anatomical distribution of the neurons. We provide concluding remarks in § 6.

## 2 Methodological framework

In this section, we describe the methodological components of the proposed vR-FCD (van Rossum-Functional Community Detection) framework (Fig. 1):

i. Starting from neural recordings of spike times (panels (a), (b)), we compute the pairwise van Rossum distances (§ A.1) between all neurons to construct a distance matrix (panel (c)).
ii. This matrix is then transformed into a weighted functional similarity network (§ A.2) using a *min-max* transformation, where nodes represent neurons and edge weights quantify similarity (panels (d), (e))
iii. Functional neural assemblies are then identified using the Louvain community detection (§ A.3) (panel (f)).
iv. To assess the performance of the vR-FCD framework, we compare the detected communities against the ground truth using the *adjusted Rand Index (ARI)* [22].

**Figure 1:**
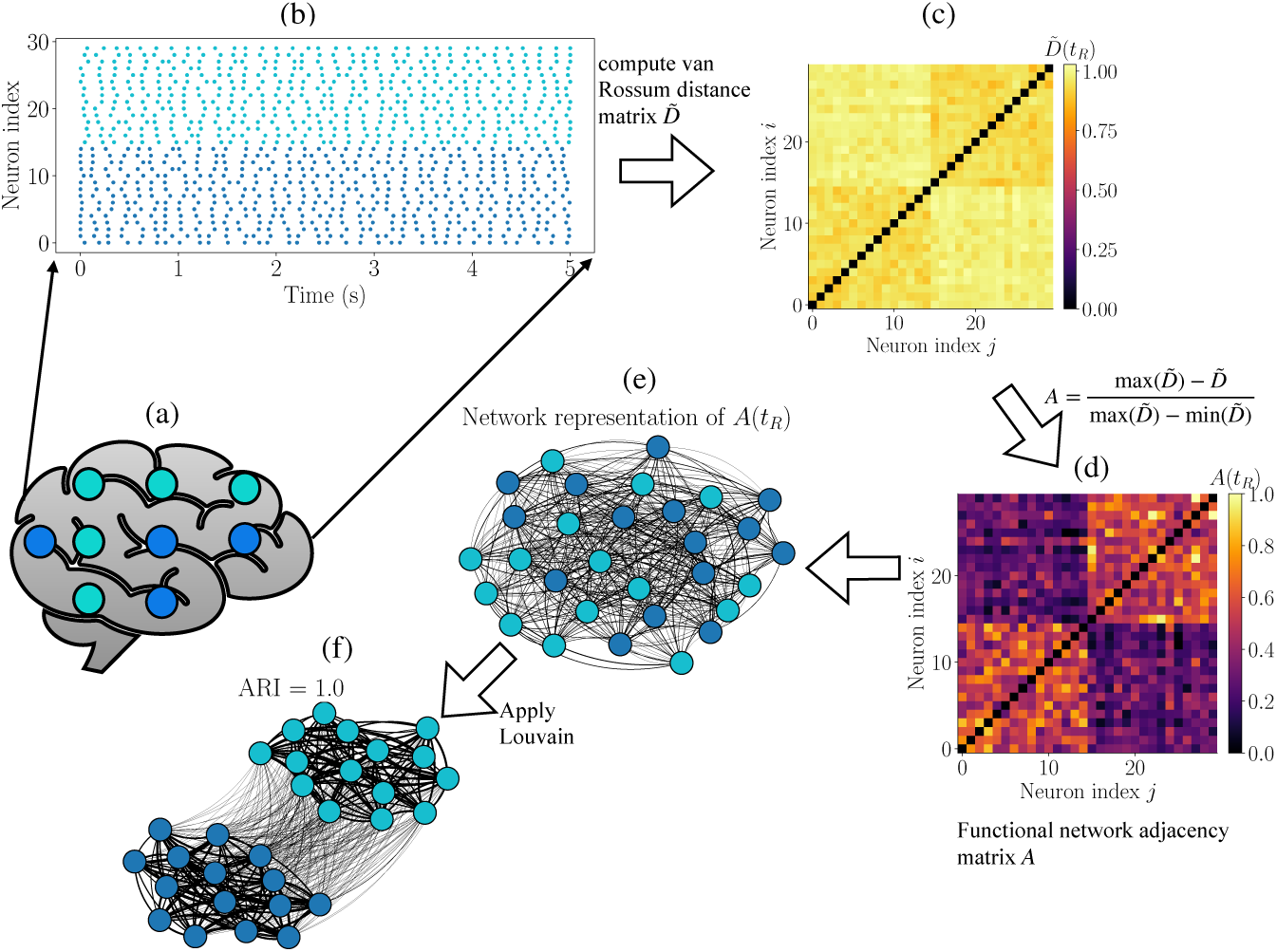
Schematic representation of the vR-FCD framework. (a) Schematic of a neural population. (b) Example spike raster with two temporally structured spike trains. (c) Pairwise van Rossum distance matrix *D̃*. (d) Corresponding functional similarity matrix *A* generated by applying the min-max normalisation to *D̃* and subtracting the values from 1. This highlights the block structure where neurons with similar temporal dynamics exhibit stringer connections. (e) Weighted functional network constructed from *A*, where nodes represent neurons and the weighted edges (thicker edges correspond to higher values) encode the temporal similarity. (f) Communities identified using the Louvain algorithm, recovering the underlying assembly structure with the ARI value displayed.

For a more detailed information on the measures, see Appendix A. We validate the framework on synthetic datasets and on networks of conductance-based LIF neurons with planted communities, before applying it to Neuropixels recordings. For the experimental data, where ground truth is unavailable, we device a separate robustness analysis and examine how the detected functional assemblies correspond to the anatomical distribution of neurons in the brain. The rationale behind using the min-max transformation to generate the similarity matrix is that it is parameter-free, preserves the full range of pairwise similarities, adapts automatically to the data, and consistently provides the most robust community recovery for the synthetic, simulated, and the Neuropixels dataset. All simulations and analyses are carried out in `Python` version 3.13.5 using its usual scientific suite: `numpy`, `scipy`, `pandas`, and `matplotlib`. More details about the `Python` packages and modules used are provided in Appendix B.

## 3 Application to synthetic data

We begin by evaluating the vR-FCD framework on the synthetic spike-train data with a known community structure. The neurons are divided into *B* blocks of equal size (*N_B_*), giving a total population of *N* = *B × N_B_*neurons. Neurons within the same block have the same firing rate characterised by a common inter-spike interval (ISI), defined as the time between successive spikes, while neurons in different blocks are separated by phase offsets. The phase shift for the *b*^th^ block relative to the first block is given by Φ*_b_* = *ϕ_b_* × ISI. There is also a jitter parameter *σ* which introduces variability in spike timing and controls the noise level within each block. Figure 2 shows a schematic example of a synthetic raster with three planted blocks illustrating the effects of these control parameters. For our simulations using the synthetic data, we keep *σ* ∈ [0, 0.4] and *ϕ_b_* [0, <u>^1^</u>] as the fractions of the ISI. Next, we test the performance of the vR-FCD framework on different control parameters of the synthetic data.

**Figure 2:**
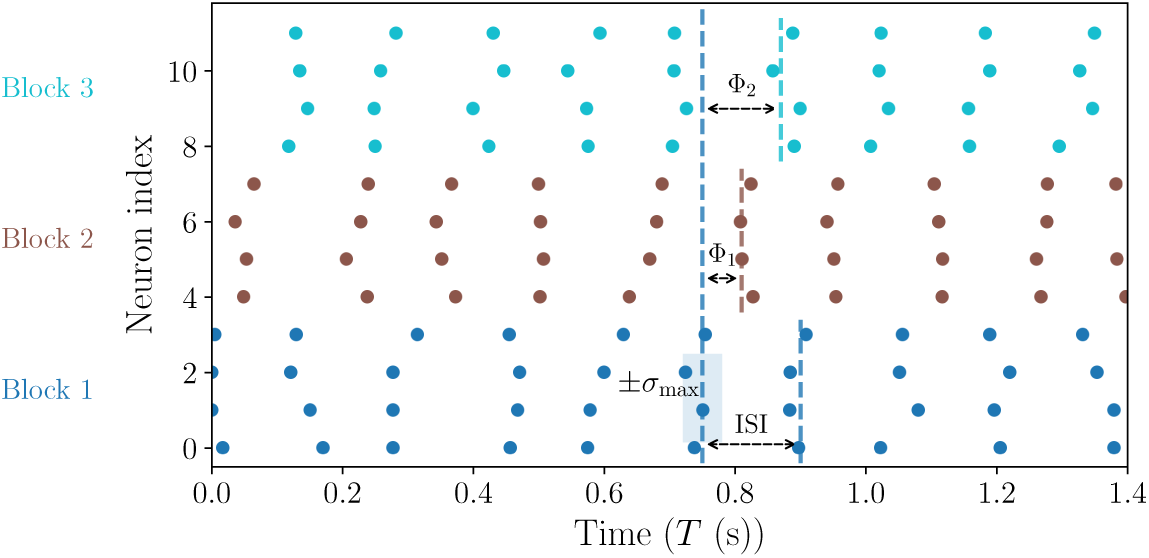
Schematic visualisation of the synthetic spike-train dataset. Neurons are divided into three blocks, each characterised by a common ISI, phase offsets *ϕ_b_* between blocks, and jitter (noise) label *σ*. The parameters Φ_1_ and Φ_2_ control the relative timing shifts between the spike trains of different blocks, while *σ* introduces variability around the mean periodic firing times.

### 3.1 Assessing the minimum recording duration *T* required for a reliable performance of the vR-FCD framework

We begin by assessing how much data is required for the vR-FCD framework to reliably recover functional communities. This is done for moderate jitter *σ* = 0.2 (Fig. 3) and high jitter *σ* = 0.4 (Fig. 4). Increasing the duration *T* improves performance by providing more spike events and more reliable estimates of spike-train similarity. This can be seen in the progressive appearance of clearer block structures in the similarity matrix *A* (within-community similarities become increasingly distinguishable from between-community similarities. For *σ* = 0.2, the underlying underlying functional structure is strong enough such that the vR-FCD framework achieves perfect recovery with ARI = 1 even for *T* = 5 s. For *σ* = 0.4, the functional structure is obscured at *T* = 5 s and the recovery is poor with ARI = 0.148. Increasing *T* to 50 100 s restores the community structure and again gives perfect recovery with ARI = 1. These demonstrate that *T* and *σ* interact strongly. Higher levels of jitter require longer recordings for the vR-FCD framework to reliably infer functional assemblies. For the rest of the simulations we use *T* = 100 s to ensure communities can be reconstructed reliably at all jitter levels.

**Figure 3:**
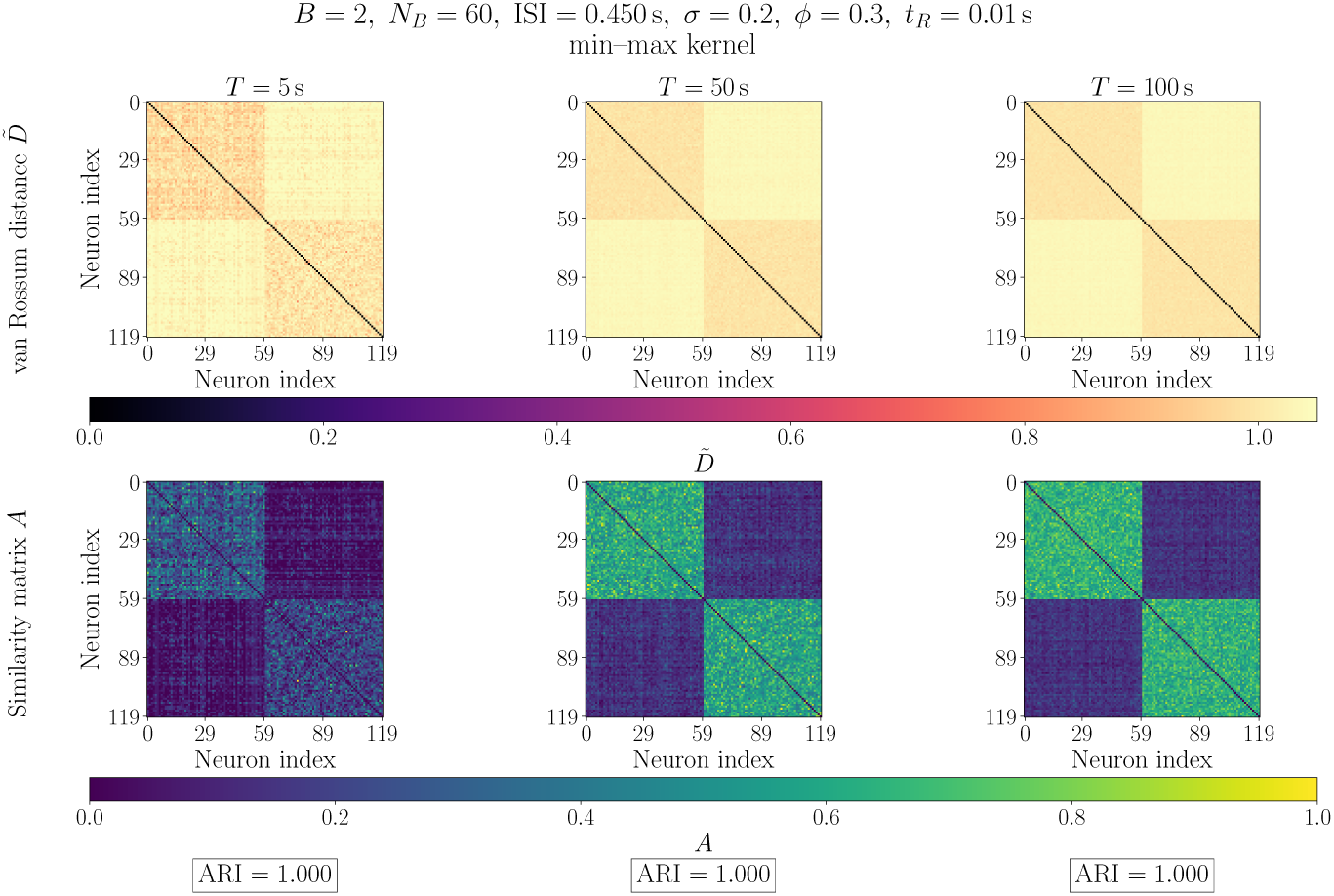
Effect of recording duration *T* on the vR-FCD framework. The synthetic dataset consists of two equally sized planted communities with jitter *σ* = 0.2. The first row shows *D̃* and the second row shows the corresponding *A* matrices. As *T* increases, the similarity estimates become more stable and the underlying block structure becomes increasingly distinct. Despite the greater variability at short durations, the separation between the within-block and between-block similarities is sufficient for the vR-FCD framework to recover the planted communities perfectly, giving us ARI = 1 for all three *T* values.

**Figure 4:**
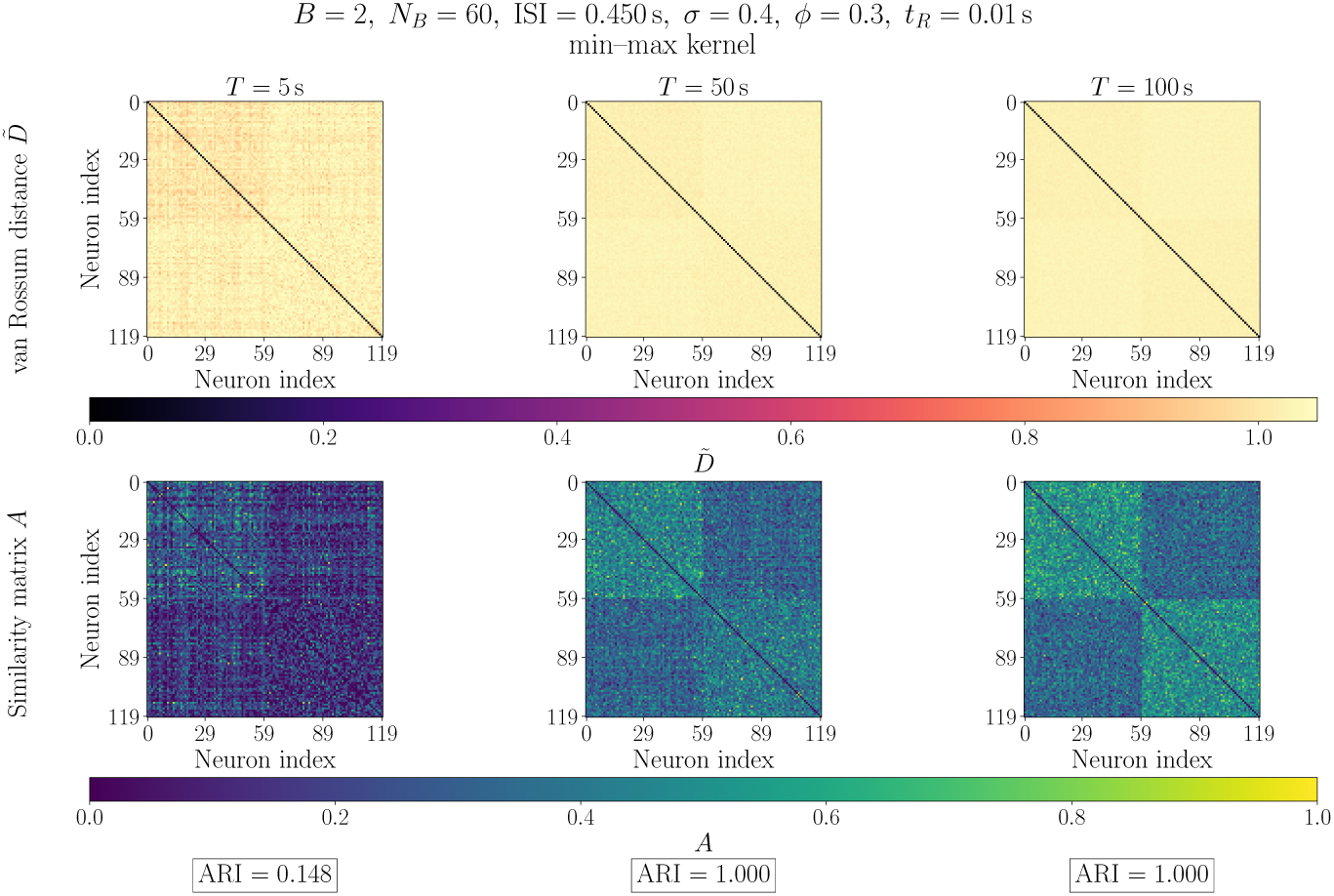
Effect of recording duration *T* on the vR-FCD framework. The synthetic dataset consists of two equally sized planted communities with *σ* = 0.4. The first row shows *D̃* and the second row shows the corresponding *A* matrices. Compared with lower jitter case in Fig. 3, higher jitter reduces the contrast between communities, making recovery more difficult at smaller *T* values. As *T* increases, the similarity estimates become more stable, enabling perfect recovery for *T* = 50 and 100 s but not for *T* = 5 s.

### 3.2 Effect of phase separation *ϕ* and jitter *σ* on accuracy value ARI

We next assess the performance of the vR-FCD framework as a function of the phase separation *ϕ* (*ϕ* = *ϕ*_1_ for *B* = 2 and *ϕ*_1_ = *ϕ*, *ϕ*_2_ = 2*ϕ*_1_ for *B* = 3) at different jitter levels *σ*, see Fig. 5. Recovery depends on whether the phase separation between the blocks remain sufficiently large relative to the spike-time variability introduced by the jitter. For a small phase separation *ϕ* the blocks are almost indistinguishable, as such the recovery performance is poor. As *ϕ* increases, the functional similarity structure becomes progressively clearer and the vR-FCD framework gives perfect recovery. The recovery is poor for *ϕ ∽*1 (*B* = 2) and *ϕ ∽* 0.5 (*B* = 3), because a phase shift of one full ISI is equivalent to *ϕ* = 0, making the two blocks overlap. For larger *σ*, the range of *ϕ* yielding near-perfect recovery becomes progressively narrower. Longer *T* partially compensate for the increased *σ* by improving recovery across the entire range of *ϕ*.

**Figure 5:**
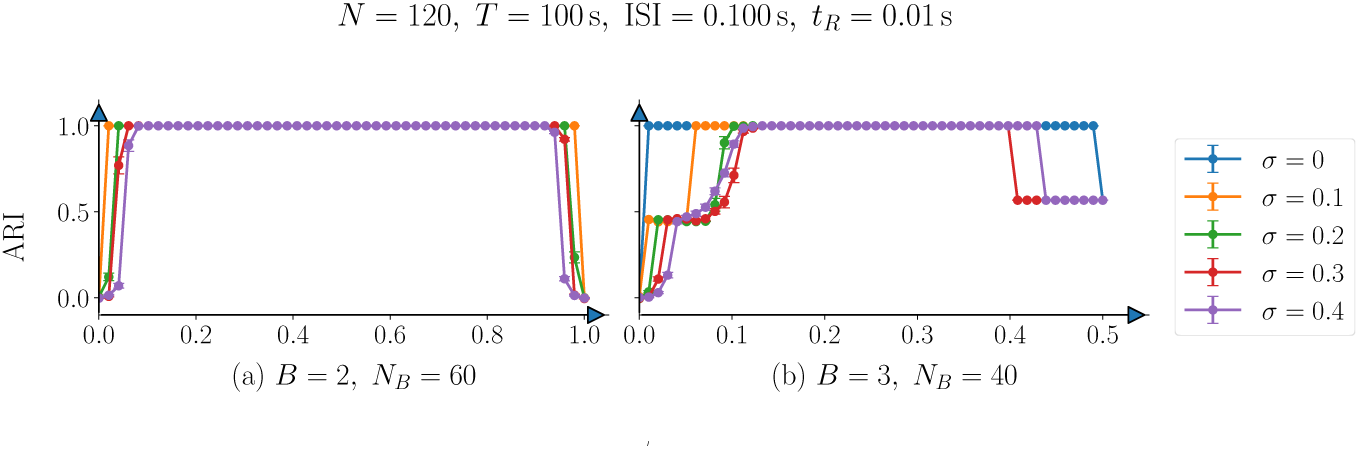
ARI as a function of *ϕ* for different *σ* levels with *T* = 100 s. Mean ARI ± standard error is shown for *B* = 2 (left) and *B* = 3 (right), with *N* = 120, ISI = 0.1 s, and *t_R_* = 0.01 s, and *N_B_* = 60 (*B* = 2) or 40 (*B* = 3). Curves correspond to *σ* = 0, 0.1, 0.2, 0.3, 0.4. ARI increases with phase separation before decreasing at large *ϕ* because of the circular phase geometry, with the effect being more pronounced for *B* = 3 owing to partial merging of communities. Increasing *σ* reduces recovery, whereas longer *T* values produce transitions and more robust community detection.

### 3.3 Effect of network size *N* and varying splits on the accuracy value ARI

Determining the operational limits of the vR-FCD framework requires identifying the lower bound of network size (*N*) at which functional assemblies remain detectable. To identify this critical threshold, we systematically varied *N* ∈ [10, 150] for *B* = 2 and *N* ∈ [10, 300] for *B* = 3 across different community split ratios (Fig. 6) for a weak phase separation *ϕ* = 0.1 and substantial jitter *σ* = 0.4. Community recovery generally improves as *N* increases, indicating that larger neural populations provide more reliable estimates of the underlying structure. For *B* = 2, our framework achieves near-perfect recovery across most *N* values, with performance degrading only for very small networks and highly imbalanced splits. For *B* = 3, larger *N* is required to achieve comparable recovery, reflecting a critical minimum community size *N_B_*. For equal sized communities, maintaining the same *N_B_* when increasing from *B* = 2 to *B* = 3 requires approximately 50% more neurons in total. Consequently, smaller or strongly imbalanced communities reach this effective threshold only at a larger *N*, making them more susceptible to merging during community detection giving lower ARI values. Beyond the ranges reported here, increasing *N* produces no appreciable changes in the observed behaviour.

**Figure 6:**
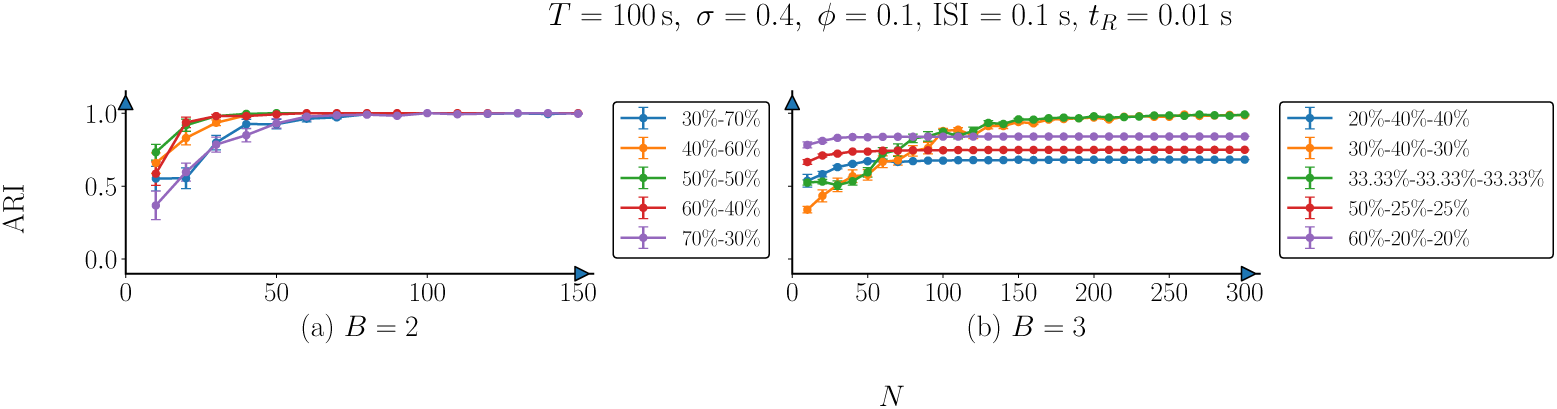
ARI as a function of *N* under high *σ* and weak *ϕ*. Mean ARI standard error is shown for *B* = 2 (left) and *B* = 3 (right), with *σ* = 0.4, *ϕ* = 0.1, and ISI = 0.1 s. We see a near-perfect ARI for *B* = 2 for *N >* 100 approximately (otherwise finite-size effect leads to degradation in ARI). The performance depends on the split balance for *B* = 3, with the imbalanced ones showing moderate ARI.

### 3.4 Effect of the time-scale parameter *t_R_* and jitter *σ* on the accuracy value ARI

A key question is to examine whether community recovery is robust to the choice of the van Rossum time scale parameter *t_R_*, which controls the temporal resolution of the spike-train comparison, i.e, smaller *t_R_*makes the distance more sensitive to fine spike-timing differences, whereas larger *t_R_* smooths activity over a broader time scale. We examine the recovery performance of the vR-FCD framework across different *t_R_* and jitter *σ* (Fig. 7). We let *t_R_* ∈ [0.001, 0.1] s, ISI = 0.1 s, and *ϕ* = 0.1 fixed, meaning the temporal separation between neighbouring blocks is 0.1 × 0.1 = 0.01 s. We find that the performance of the framework remains high when *t_R_* is small relative to the characteristic temporal separation between communities. It deteriorates as *t_R_* increases and temporal differences become progressively smoothed out. This is more evident in the noisy regimes, where broader kernels reduce the contrast between the blocks. The dependence on *t_R_* is weak for *B* = 2 case, indicating that the underlying functional structure remains sufficiently distinct across a broad range of *t_R_* values. Increasing *t_R_* leads to a partial merging of communities and reduction in ARI for the most noisier (*σ* = 0.4) case. For *B* = 3, the separations between blocks are smaller, creating partial overlaps between communities. A near-perfect recovery occurs for *σ* ≤ 0.1, but as *σ* increases ARI stabilises near 0.45 − 0.55. The three-block structure partially merges, but some mesoscale organisation still survives. Earlier duration analyses showed that increasing *T* further improves recovery, indicating that more data are required when distinguishing a larger number of communities with relatively small phase separations. A successful community recovery occurs when *t_R_* ≲ *ϕ* × ISI, while recovery deteriorates once *t_R_*∼ ISI.

**Figure 7:**
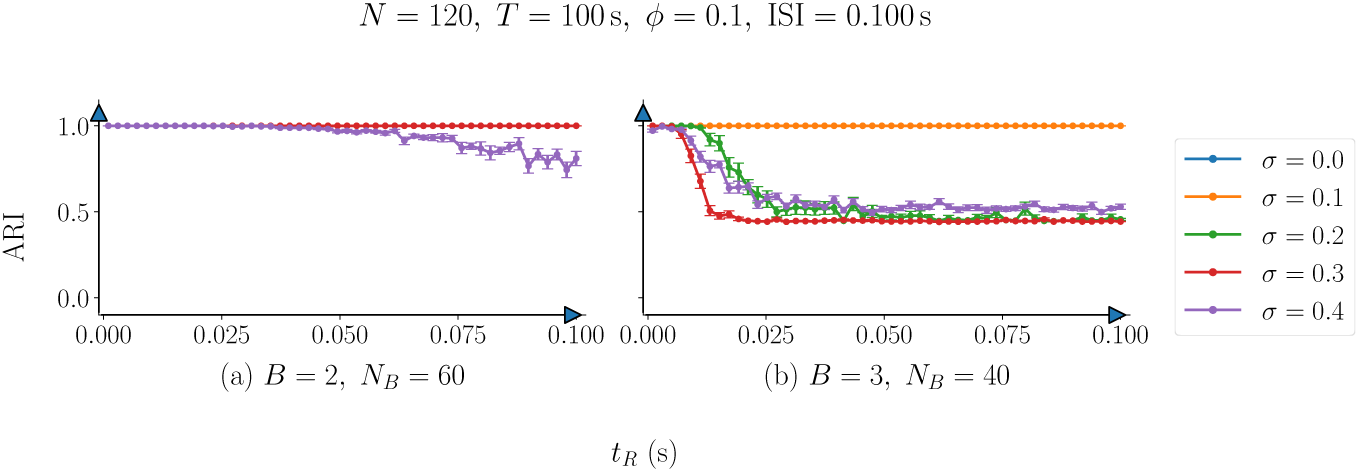
ARI as a function of the time scale parameter *t_R_* with *ϕ* = 0.1, ISI = 0.1 s, and *T* = 100 s. The temporal separation between neighbouring communities is *ϕ* ISI = 0.01 s. Increasing *t_R_* reduces ARI, particularly at high *σ*, as broader temporal kernels blur distinctions between communities. This effect is stronger for *B* = 3, where community overlap limits ARI to approximately 0.45 − 0.55 at large *t_R_*. Reliable recovery occurs when *t_R_* ≲ *ϕ* × ISI and deteriorates once *t_R_*becomes comparable to the ISI.

### 3.5 Effect of varying the phase separations *ϕ*_1_ and *ϕ*_2_ on the accuracy value ARI for three-blocked planted structure

A remaining question is whether the performance observed for *B* = 3 depends on the imposed phase geometry (with phase differences *ϕ*_1_ and *ϕ*_2_), or persists when the three communities are positioned independently around the phase circle with jitter values *σ* = 0.2 and *σ* = 0.4 (Fig. 8). In our previous experiments a single parameter *ϕ* was used to prescribe the relative phase offsets between the three planted blocks. Here we relax this constraint by taking block 1 as the reference and defining *ϕ*_1_ as the phase offset between blocks 1 and 2, and *ϕ*_2_ as the phase offset between blocks 1 and 3. We vary (*ϕ*_1_*, ϕ*_2_) over a 100 × 100 grid retaining only points which satisfying the inequality

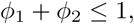

as indicated by the red boundary in Fig. 8. We run 20 independent simulations on each grid point and take the mean ARI. Since the phase offsets lie on a periodic domain, the effective separation between any two blocks is determined by their shortest distance around the phase circle. Hence, the three-community recovery is governed by the pairwise separations among all three blocks, including the separation between the second and the third blocks.

**Figure 8:**
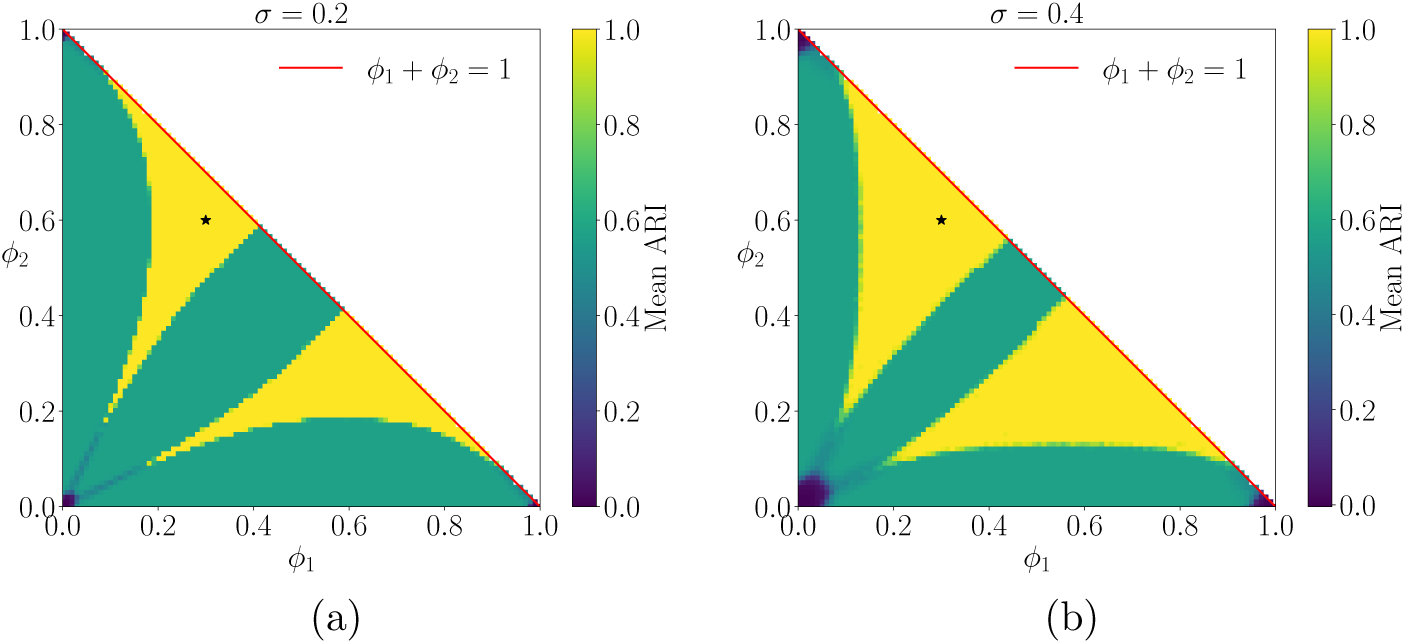
Mean ARI computed over a 100 × 100 grid of phase pairs (*ϕ*_1_*, ϕ*_2_) satisfying *ϕ*_1_ + *ϕ*_2_ ≤ 1 (red boundary). The phases are (0*, ϕ*_1_*, ϕ*_2_) with *B* = 3, *N_B_* = 40, ISI = 0.1 s, *T* = 100 s, and *t_R_* = 0.01 s. Panels (a) and (b) correspond to *σ*= 0.2 and *σ*= 0.4, respectively. High ARI (yellow colour) is obtained when the communities are well separated in phase, whereas low ARI occurs when two or more phases overlap because of circular wrapping. The black star denotes the well-separated configuration (*ϕ*_1_*, ϕ*_2_) = (0.3, 0.6). Increasing jitter broadens the low-ARI regions and smooths the transition between successful and failed recovery.

Recovery is successful when all three separations are large enough and fails whenever one pair of blocks becomes phase aligned nearly. For example, the point (*ϕ*_1_, *ϕ*_2_) = (0.3, 0.6), marked by the black star, corresponds to a good separation between the blocks and ultimately a near-perfect recovery. At (*ϕ*_1_, *ϕ*_2_) = (0, 0) however, ARI ≈ 0 because the phases are identical. Similar reductions in recovery occur near (*ϕ*_1_*, ϕ*_2_) ≈ (0, 1) and (*ϕ*_1_*, ϕ*_2_) ≈ (1, 0). High-performance regions occur at the centre of the triangular domain, where all three blocks are well separated around the circle, whereas low-performance blocks appear near the corners with partial recovery along the *ϕ*_1_ ≈ *ϕ*_2_ line and *ϕ*_1_ or *ϕ*_2_ ≈ 0.

We also observe an interesting effect of jitter. Increasing *σ* from 0.2 to 0.4 does not produce a simple uniform plummet in performance. Some well-separated regions become broader and remain in the high-recovery yellow regime, suggesting that the vR-FCD framework is still recovering the planted communities when the phases are well separated. We see that the boundary between perfect recovery to partial recovery broadens and smoothens. However, for *σ* = 0.4 there are also larger dark regions where two or more phases are close. This means jitter amplifies the dependence on phase geometry, i.e, well-separated configurations can still be recovered robustly, whereas poorly separated configurations fail more. For example when (*ϕ*_1_*, ϕ*_2_) = (0.14, 0.6), the ARI increases from approximately 0.567 for *σ* = 0.2 to 1 for *σ* = 0.4. In this case, the increased jitter alters the pairwise spike-train similarity structure such that communities that are only partially resolved at lower jitter become more clearly separated. We conclude that the jitter *σ* controls how sharply the vR-FCD framework transitions between failure, partial recovery and perfect recovery of the planted communities. These results helps us build an all-round benchmark around our methodology.

## 4 Application of the vR-FCD framework to a stochastic block model (SBM) network of Leaky Integrate and Fire (LIF) neuron model

Having established the performance of the vR-FCD framework under controlled perturbations, we employ the framework to spike trains generated by a network of spiking neurons. We generated a spike raster from simulating a network of conductance-based Leaky Integrate and Fire (LIF) neurons, see Appendix D. The network is a *stochastic block model* [23] (SBM) of *N* neurons consisting of *n_C_* communities (or ‘blocks’ to draw analogy with our synthetic spike train dataset from the previous section) which we plant beforehand and construct the adjacency matrix M accordingly. Here, we consider two planted communities (*n_C_* = 2), but present results for *n_C_* = 3 in Appendix F. The connection probabilities are given by *p*_in_, and *p*_out_. The network structure is governed by two key probabilities: *p*_in_, which sets the probability of a directed connection between any two neurons within the same community, and *p*_out_, which sets the probability of a directed connection between neurons in different communities. These prepare a ground-truth structural network of the neurons which is utilised later to validate the community detection from the similarity network, i.e, the functional network of the simulated spike train from this network of LIF neurons. Within each cluster, we set 80% of the neurons to be excitatory, with the remaining 20% being inhibitory.

When a neuron spikes, it modifies the synaptic conductance of its connected neighbours according to whether it is excitatory or inhibitory. We denote the corresponding coupling strengths by *g_E_* and *g_I_*. Since the balance between excitation and inhibition can influence the community recovery, we later vary *g_E_* and *g_I_* to assess its effect on the community recovery.

The planted communities are defined by the SBM adjacency matrix, where each neuron also receives an external *Poisson* drive that modulates its excitatory input. In later experiments, we vary the drive parameters between clusters to examine how heterogeneous external input alters the spike-train similarity structure and community recovery without changing the underlying structural modularity. Simulations are run for a duration of *T* s. Further details of the Poisson input and synaptic dynamics are provided in Appendix D. For our spiking neuron model, we adopt the LIF neurons with parameters fit to match the spiking activity of Hodgkin–Huxley [24] type descriptions of cortical neurons [25, 26]. The parameter set fixed in this paper is given in Tab S1. For the simulation, we integrate the ODE using the second-order Runge Kutta method (RK2) with Δ*t* = 0.0001 s over a duration of *T* = 20 s. An outline of the network simulation, generation of the corresponding spike-activity raster, and application of our vR-FCD framework on the simulated spike raster is provided as a pseudocode in Appendix E.

### 4.1 Case study with one simulation run

To provide an interpretable benchmark before the full parameter sweep, we first analyse a representative simulation in which the planted structural communities are clearly defined and their corresponding spiking activity can be inspected directly. The network has *N* = 100 neurons with *n_C_* = 2 equally sized blocks of 50 neurons each, along with within-cluster probability *p*_in_ = 0.7 (dense) and the between-cluster probability *p*_out_ = 0.01 (sparse) (see Fig. 9 (a) for a representation of the SBM adjacency matrix and (b) for the corresponding directed network representation). The goal is to mimic distinct neural assemblies that are internally well-connected but loosely connected to each other. We set *r_C_* = 100 Hz and *f_C_* = 0.2 such that the network exhibits sustained irregular spiking with a mean population firing rate of approximately 30 Hz and coefficient of variation (CV) of the ISI distribution of roughly 1. The nodes in the network are coloured by cluster membership, illustrating a two-community topology. The network is simulated for 20 s and the spike times of each neuron are recorded. Figure 9 (c) shows the raster plot, with the neurons coloured by cluster similar to the network. For visual clarity, we show representative subset of the time course only.

**Figure 9:**
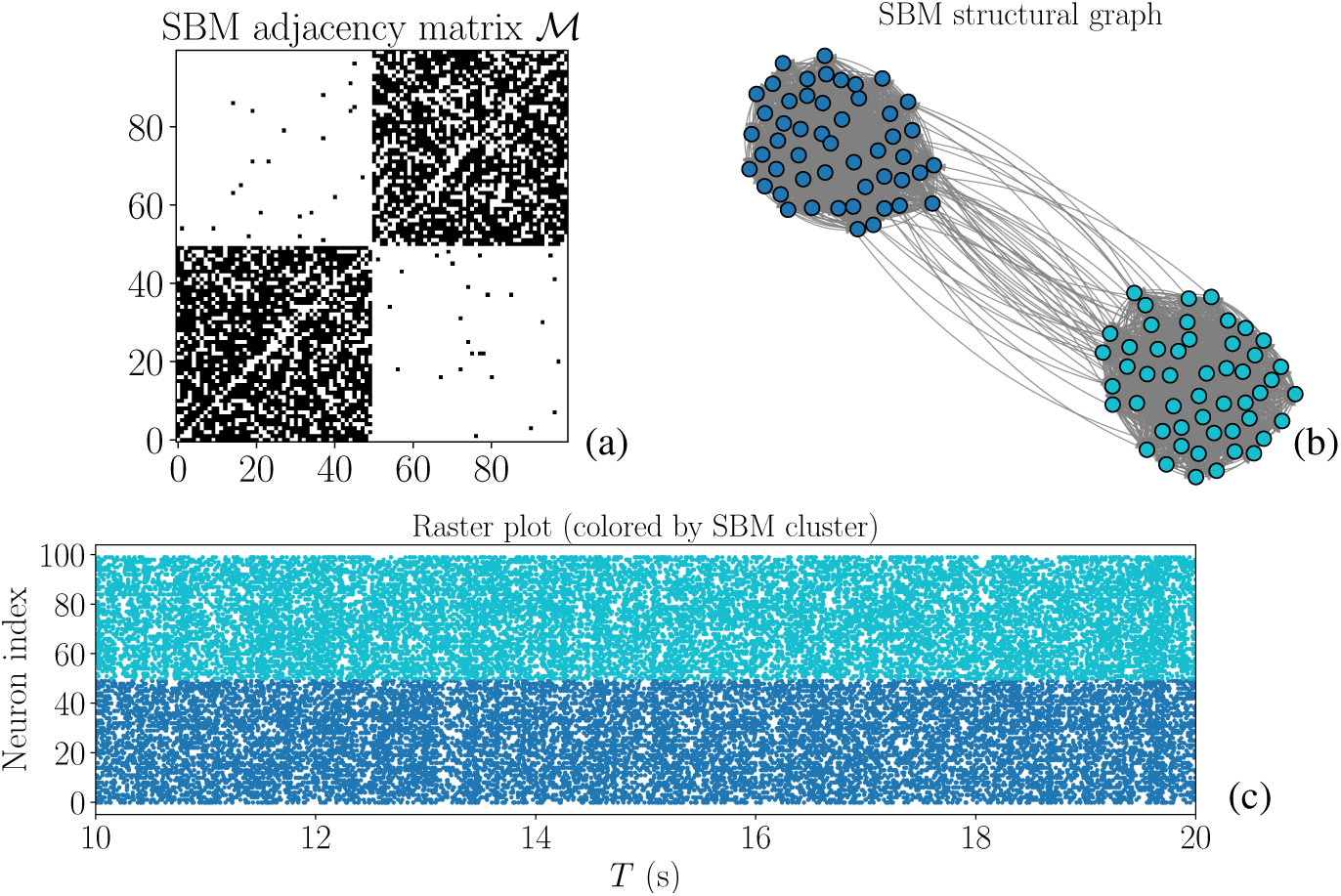
Stochastic Block Model (SBM) network of a Leaky Integrate and Fire (LIF) neurons model and the resulting spike activity. (a) Binary asymmetric SBM adjacency matrix M with dense within-cluster (*p*_in_ = 0.7) and sparse between-cluster (*p*_out_ = 0.01) connectivity. (b) Corresponding directed structured network. The simulation consists of *N* = 100 neurons divided equally into two communities with *p_E_* = 0.8. (c) Simulated spike raster under external Poisson drive (*r_C_* = 100 Hz, scaled by *f_C_* = 0.2) showing the last *T* = 10 s of the 20 s simulation. The network exhibits a sustained, irregular activity with distinct but overlapping temporal patterns across the two clusters, providing a controlled benchmark for evaluating the vR-FCD framework.

Using the generated spiking data, we apply the vR-FCD framework (with *t_R_* = 0.005 s) to assess community recovery accuracy. On comparing the detected communities from the spike-train similarity network with the planted SBM partition, we obtain an ARI of 0.8824 (Fig. 10). The recovery is high but not perfect because structural co-membership does not guarantee identical spike dynamics. External drive, inhibitory/excitatory interactions, and finite recording duration introduce within-block variability. This leads to some neurons be functionally more similar to neurons outside their planted block. The resulting mismatch is therefore expected from the dynamics of the LIF network. The same issue is likely to be even more pronounced in experimental spike recordings where functional activity need not map one-to-one onto anatomical or structural connectivity. Note that the framework easily scales up to more clusters, see Appendix D.

**Figure 10:**
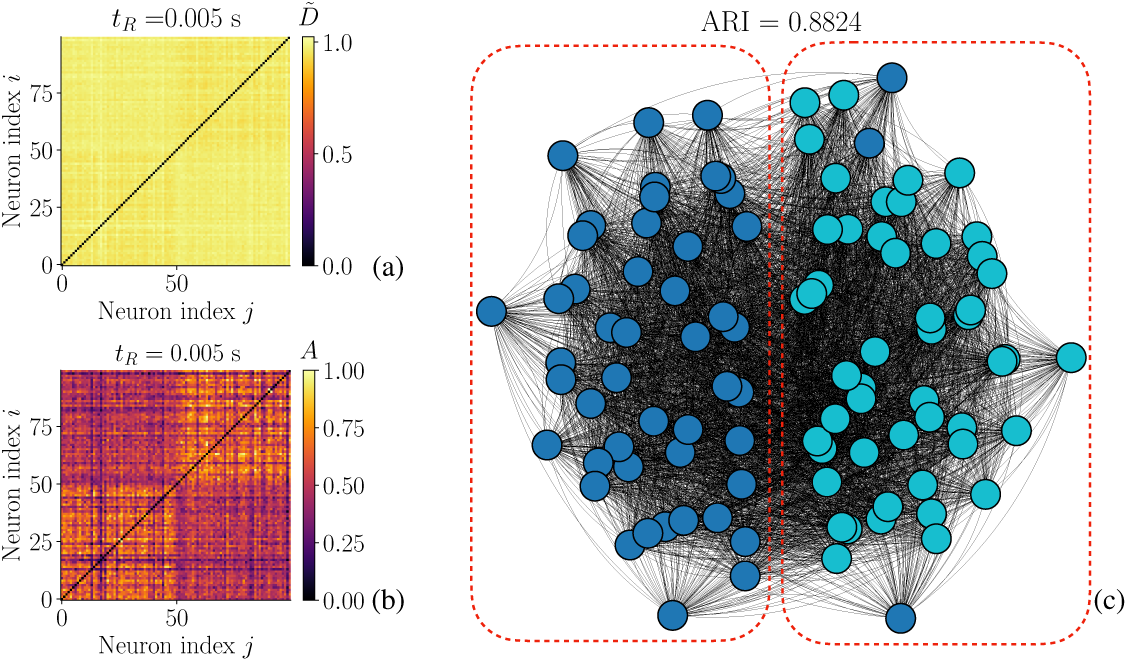
Community detection in the LIF neuron network via the vR-FCD framework. **(a)** Pairwise van Rossum distance matrix *D̃* (*t_R_* = 0.005 s) computed from the spike raster in Fig. 9 (c). (b) Corresponding similarity matrix *A*, revealing a clearer two-community structure. (c) Communities detected by the vR-FCD framework, recovering clusters of sizes 53 and 47 (ARI = 0.8824), close to the planted SBM partition. Although *D̃* exhibits weak block structure, the similarity transformation enhances community separability.

### 4.2 Performance of the vR-FCD framework across two-dimensional parameter spaces

In this section we investigate how network structure, synaptic interactions, and external drive affect community recovery in the LIF network (Fig. 11). Unless stated otherwise, we keep the parameter values unchanged from those introduced in the previous sections. We start by varying the within and between cluster connection probabilities, *p*_in_ and *p*_out_, respectively (Fig. 11 (a)). The results reveal a clear transition between successful and unsuccessful recovery that is governed by the balance between *p*_in_ and *p*_out_. High ARI values occur when *p*_in_ is large and *p*_out_ is small, indicating that the functional spike-train dynamics retain the modular organisation present in the structural network. Biologically, this regime corresponds to a strongly segregated modular architecture, characterised by dense within-community and sparse between-community connectivity, a widely observed organisational principle of structural and functional brain networks [27]. As the distinction between *p*_in_ and *p*_out_ decreases (top left corner of the surface plot), the communities become progressively mixed and recovery deteriorates. As expected, the firing rate remains nearly constant (range 30 − 31 Hz) across the parameter plane.

**Figure 11:**
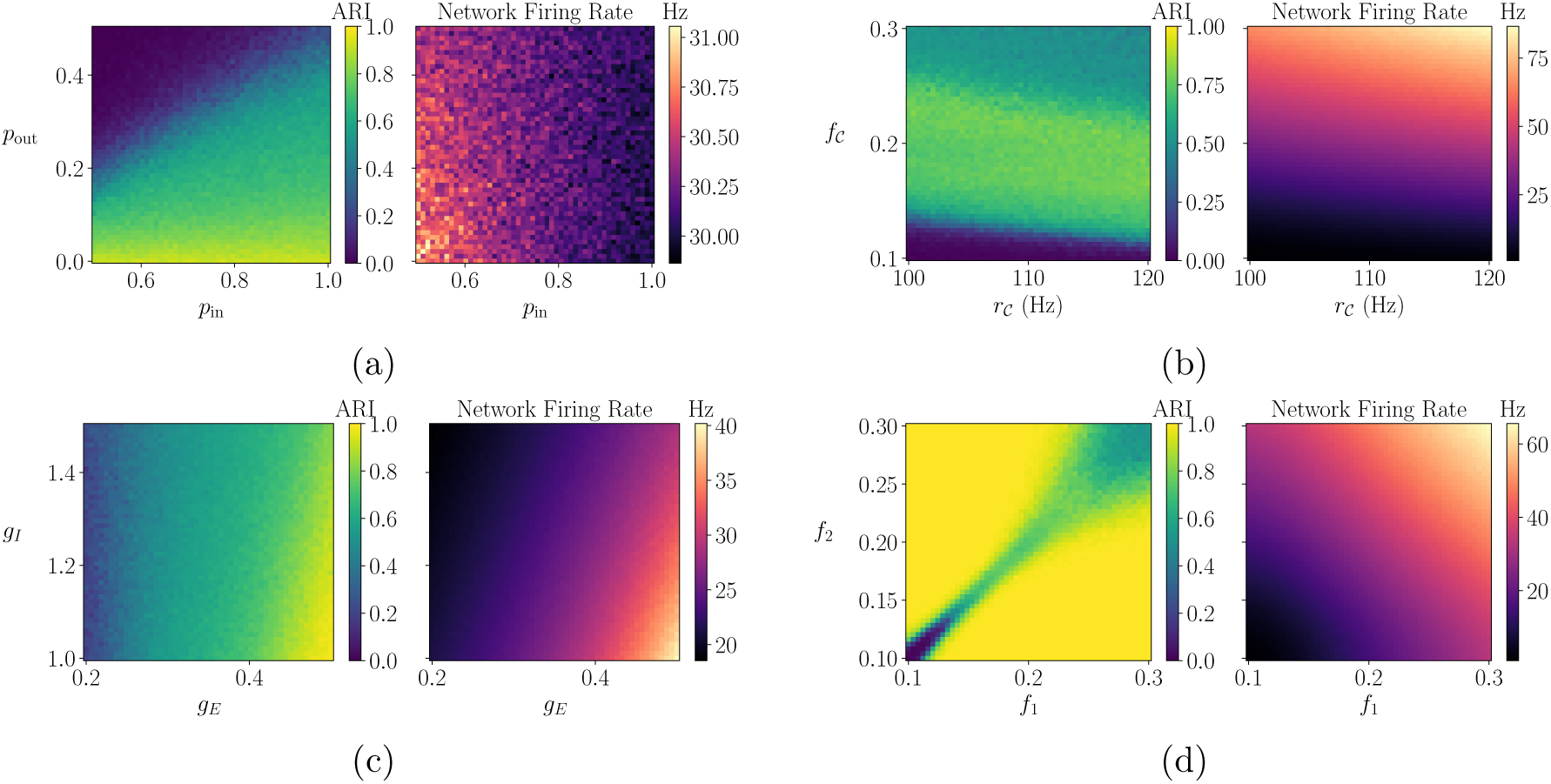
Two-dimensional parameter sweeps showing the mean ARI (left) and the firing rate (right) for the two-cluster SBM. Simulations use *N* = 100, *N_B_* = 50, *t_R_* = 0.005 s, *p_E_* = 0.8, *T* = 20 s, Δ*t* = 0.0001 s, and parameters from Tab. S1, with results averaged over 20 runs. Panels show the effects of varying (a) *p*_in_ and *p*_out_ with *g_E_* = 0.4, *g_I_* = 1 and *r_C_* = 100 Hz, *f_C_* = 0.2 on both clusters, (b) *r_C_* and *f_C_* with *g_E_* = 0.4, *g_I_* = 1, *p*_in_ = 0.7, and *p*_out_ = 0.1, (c) *g_E_* and *g_I_* with *p*_in_ = 0.7, and *p*_out_ = 0.1, and *r* = 100 Hz, *f* = 0.2 on both clusters, and (d) *f*_1_, *f*_2_ with *r_C_* = 100 Hz, *g_E_* = 0.4, *g_I_* = 1, *p*_in_ = 0.7, and *p*_out_ = 0.1. Community recovery is the strongest for highly modular networks, sufficient excitation, and distinct external inputs, but deteriorates as the two communities become dynamically indistinguishable.

Having established a baseline recovery of 88% for the fixed modular network, we next investigate how external input affects the vR-FCD performance (Fig. 11 (b)). We vary the rate and amplitude of the external Poisson drive, *r_C_* ∈ [100, 120] Hz and *f_C_* ∈ [0.1, 0.3], while keeping the underlying SBM connectivity unchanged. Recovery is poor for *f_C_* ≈ 0.1, highest for intermediate *f_C_* ≈ 0.15 − 0.25, and decreases again towards *f_C_* ≈ 0.3. The firing rate (right) shows that the firing rate increases with *f_C_* with almost 75 Hz for *f_C_* = 0.3. The reduction in ARI at different drives may occur because *t_R_* = 0.005 s is kept fixed even though the firing rate, and therefore the typical ISI, changes. We examine this further in § 4.4 by varying *t_R_* for different external drive amplitudes.

The previous results suggest that, beyond the underlying modular structure and external drive, the network dynamics may also influence how clearly the communities can be recovered. To test this we vary the synaptic strengths of the excitatory and inhibitory neurons *g_E_* ∈ [0.2, 0.5] and *g_I_* ∈ [1, 1.5], respectively (Fig. 11 (c)). Stronger excitation amplifies the activity patterns associated with the underlying SBM communities, hence ARI increases with increases in *g_E_* for *p*_in_ *> p*_out_. Neurons in the same cluster begin to share more similar temporal activity, and the van Rossum distance better reflects the SBM blocks. The dependence on *g_I_* on the other hand is less pronounced. Strong inhibitory synapse suppresses spiking which can reduce the performance of recovering the within-cluster temporal activity pattern, hence increasing *g_I_* weakens recovery. We also note that with increasing *g_E_*, the firing rate increases, indicating that stronger excitatory coupling produces higher overall spiking activity. While firing rates increase with *g_E_* and decrease with *g_I_*, the performance cannot be explained by firing rate alone. Rather, these results suggest that our framework is most successful when the synaptic interactions enhance the modular organisation already present in the network.

To examine the effect of heterogeneous external input, we independently vary the drive amplitudes *f*_1_*, f*_2_ ∈ [0.1, 0.3] between the two clusters (Fig. 11 (d)). As expected, the ARI is the best when *f*_1_ ≠ *f*_2_. The performance drops along the diagonal *f*_1_ ≈ *f*_2_. When the two clusters receive similar external drive amplitudes, our framework performs worse. When *f*_1_ = *f*_2_, the two clusters operate in two different dynamical regimes with one cluster firing more than the other, leading to a clear distinction between the two groups. The firing rate increases smoothly with *f*_1_ and *f*_2_, reflecting increased neuronal activity under stronger external excitation, and is the highest when both drive amplitudes are large. We also tested heterogeneity in the input rates by independently varying *r*_1_*, r*_2_ ∈ [100, 120] Hz with *f_C_* = 0.2. Over this range, community recovery showed little dependence on *r*_1_ and *r*_2_, indicating that moderate differences in input rate have a much weaker effect on recovery than differences in drive amplitude.

### 4.3 Performance of the vR-FCD framework as a function of network size *N*

Here, we investigate how the performance of the vR-FCD framework scales with the network size *N* ∈ [10, 500] for five different community splits, see Fig. 12. The ARI increases fast with *N*, reaching ARI 1 by *N* 30 50, and is stable thereafter at ≈ 1. Balanced splits (50% 50%, 60% 40%, and 40% 60%) slightly outperforms the more imbalanced ones (30% 70%, and 70% 30%). The poorer performance for very small *N* reflects finite-size effects in the SBM: with only a few neurons per community, the connectivity can differ substantially from the prescribed probabilities, and some neurons may have very few connections, weakening the effective modular structure. A slight reduction in ARI is observed beyond *N* ≈ 200, which may arise from the growing number of cross-community connections and stochastic interactions as the network becomes bigger, making the functional distinction between the communities marginally less pronounced. This indicates that the performance is primarily driven by *N* rather than changes in the overall activity level.

**Figure 12:**
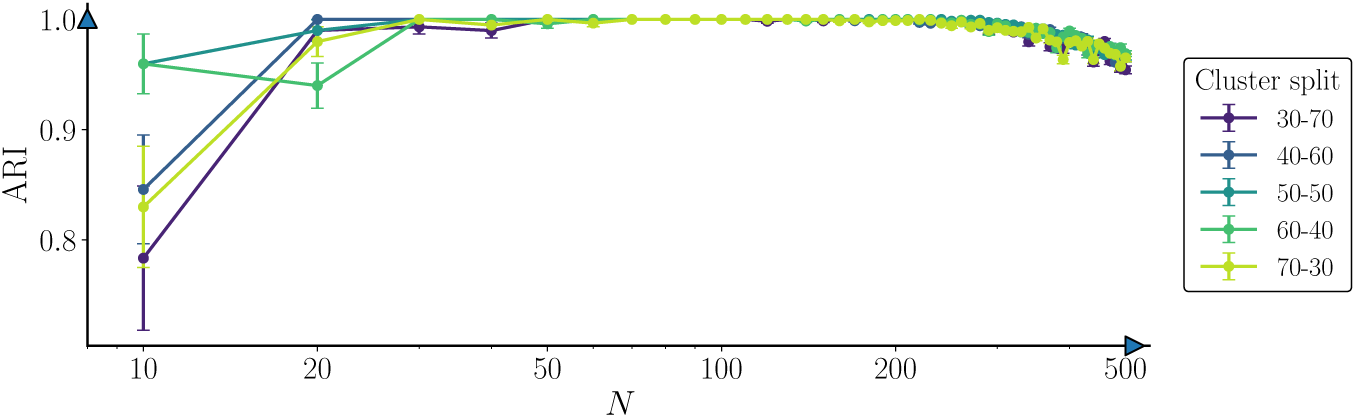
Mean ARI and network firing rate vs network size *N* for different cluster splits. Simulations use *p*_in_ = 0.8, *p*_out_ = 0.01, *g_E_* = 0.5, *g_I_* = 0.1, *r_C_* = 100 Hz, *f_C_* = 0.2, and *t_R_* = 0.005 s. Community recovery improves fast with increasing *N*, achieving ARI ≈ 1 for *N ≥* 30, with balanced community splits slightly outperforming more imbalanced ones. Results are averaged over 20 independent simulations with standard error bars.

### 4.4 Effect of the time-scale parameter *t_R_* and the drive amplitude *f_C_* on the performance of vR-FCD

To determine how the optimal *t_R_*for community recovery changes with the firing regime, we vary *t_R_*∈ [0.0001, 0.032] s across low, intermediate, and high mean firing rates of 12.07 ± 0.03 Hz, 30.36 ± 0.12 Hz, and 65.32 ± 0.26 Hz respectively (Fig. 13). The optimal *t_R_* decreases systematically as firing rate increases. This is consistent with the reduction in characteristic ISI at higher firing rates, which requires a narrower temporal kernel to resolve spike-time synchrony. The maximum ARI also tends to increase with firing rate, likely because higher firing activity provides more spike events over the same recording duration and therefore more information for estimating similarities. These indicate that *t_R_* should be chosen relative to the characteristic firing time scale of the population.

**Figure 13:**
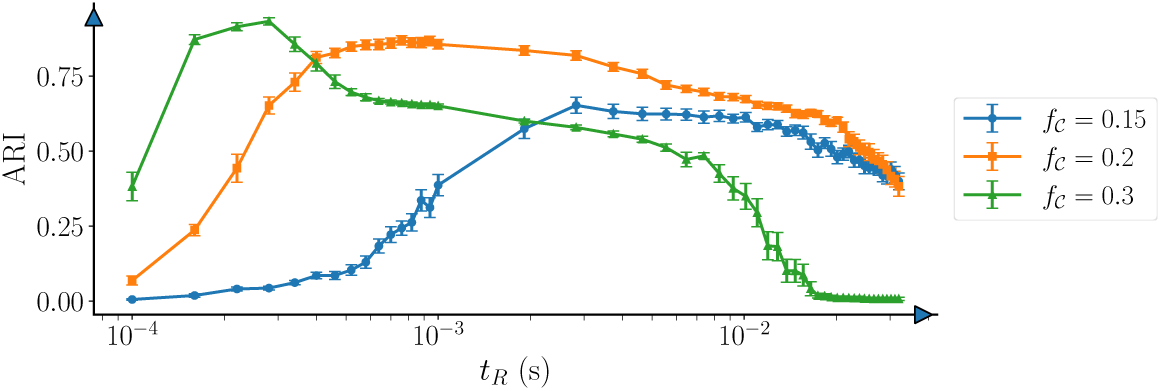
Mean ARI vs the van Rossum temporal scale *t_R_*. for different *f_C_*. Here *p*_in_ = 0.7, *p*_out_ = 0.1, *g_E_* = 0.4, *g_I_* = 0.1, *r_C_* = 100 Hz, with varying *f_C_* = 0.15, 0.2, 0.3 (with firing rates 12.07 ± 0.03 Hz, 30.36 0.12 Hz, and 65.32 0.26 Hz respectively) and *t_R_* [0.0001, 0.032] s. The optimal *t_R_* shifts towards smaller values as the firing rate increases, with each point representing the mean ARI over 20 simulations with standard error bars.

## 5 Application of the vR-FCD framework to electrophysiology data from the Allen Institute for Brain Science

With the validation in a mechanistic setting complete, we investigate whether these principles carry through to experimental recordings in the last part of this work. To this end we apply the framework to Neuropixels dataset from the Allen Institute for Brain Science [28, 29], which provided open-access recordings of *in-vivo* extracellular electrophysiology (ecephys) activity from awake mice. We focus on “spontaneous activity” from the “Functional Connectivity” dataset processed from *wild-type C57BL6/J* mice. These recordings contain spike trains from thousands of simultaneously recorded neurons across multiple brain regions, using Neuropixels probes. This offers an ideal testbed for assessing whether vR-FCD can reveal meaningful functional organisation in real data. The data is accessed through the software development kit called `AllenSDK` (`https: //allensdk.readthedocs.io/en/latest/visual_coding_neuropixels.html`). To learn more about how the experiments are designed, data are acquired, and the type of data we concentrate our focus on, see Appendix G.

After filtering we obtain 14 recording sessions comprising between 531 and 930 units each (see Tab. S2 in Appendix G for the summary of the filtered sessions, including the session metadata, number of units, probe counts, etc.). Every session corresponds to an independent Neuropixels recording from a single mouse and is analysed separately using the vR-FCD framework. The recordings are relatively homogeneous from a biological perspective, with all animals being adults (108 − 142 days old), reducing potential variability associated with developmental stage. The sessions comprise recordings from 11 male and 3 female animals and include high-density recordings obtained using 5 − 6 probes with 1696 − 2298 recording channels. To ensure robust analysis, we restrict our attention to high quality units with a signal-to-noise ratio greater than 4. Spontaneous activity is available in six presentations shared across all recording sessions (IDs: 0, 3646, 3797, 4338, 40639, 40940). These presentations have highly consistent durations across sessions, providing a common basis for comparing functional organisation between animals. The summary statistics for the stimulus durations are reported in Table. S3 (Appendix G).

### 5.1 Temporal robustness of functional communities detected using the vR-FCD framework

We begin assessment of our vR-FCD framework by performing a robustness test. Since the underlying functional communities are unknown, the goal is not validation against ground truth but rather to determine whether the framework produces stable and reliable community structure in the neural activity. We focus on the longest spontaneous recording (stimulus ID 40639; ~ 1800 s), as our earlier analyses showed that reliable recovery requires sufficiently long observations, especially when multiple communities must be distinguished. For each session, spike trains are converted into a binary raster using a bin width based on the smallest observed ISI. Across the sessions, the characteristic ISI, defined as the median of the neuron-wise median ISIs, has a median value of approximately 0.085 s (range 0.051 − 0.114 s), corresponding to a characteristic firing rate of approximately 12 Hz. Our LIF simulations show that the optimal van Rossum time scale depends on the firing regime and remains substantially shorter than the characteristic ISI (Fig. 13). For a comparable firing rate of approximately 12 Hz, the optimal *t_R_* lies on the order of a few milliseconds. We therefore use *t_R_* = 0.001 s as a conservative fine time-scale parameter, which is well below the characteristic ISI and within the range over which the LIF simulations show good recovery performance. The synthetic benchmarks showed reliable recovery for *T* = 100 s and ISI = 0.1 s, corresponding to approximately 10^3^ times the characteristic ISIs. Applying the same scaling to the Allen data would correspond to about 85 s of activity, which is substantially shorter than the 1800 s vailable in stimulus ID 40639. This therfore provides ample spike activity for estimating the functional similarity structure. After applying the vR-FCD framework, most recordings partition into two dominant communities, although a small number of sessions contain additional minor clusters consisting of only a few neurons. For example for session ID 768515987, our method generates two clusters with 36 units in both, whereas for session ID 771160300, our method gives four clusters with 71 units in the first cluster, 58 units in the second, and two units each in the third and the fourth clusters.

To asses the temporal robustness of the detected communities, we ask whether the functional partition inferred from the full spontaneous recording can be recovered from shorter portions of the same recording. In the absence of a known ground-truth partition for the Allen data, this mechanism provides a robustness test for the vR-FCD framework. If the detected communities are long-lived, partitions obtained from shorter time windows should remain similar to that obtained from the complete recording. Conversely, reduced performance can occur if the underlying functional organisation itself changes over time. We therefore treat the partition obtained from the longest spontaneous recording (stimulus ID 40639) as the reference and apply vR-FCD to fixed 10% segments and overlapping windows spanning 30%, 50%, 70% and 90% of the recording. The resulting partitions are compared with the reference using ARI. Agreement increases consistently with window length across sessions (Fig. 14 (a)). Relatively short windows retain a substantial fraction of the full-recording community structure. This suggests that the detected communities are not confined to a single time segment, but retain a substantial degree of temporal consistency across the recording, while allowing for changes in community organisation over time. At the same time, the analysis should be interpreted as a test of temporal consistency rather than ground-truth accuracy. In the absence of experimentally defined functional community labels, such within-recording reproducibility provides a practical measure of robustness.

**Figure 14:**
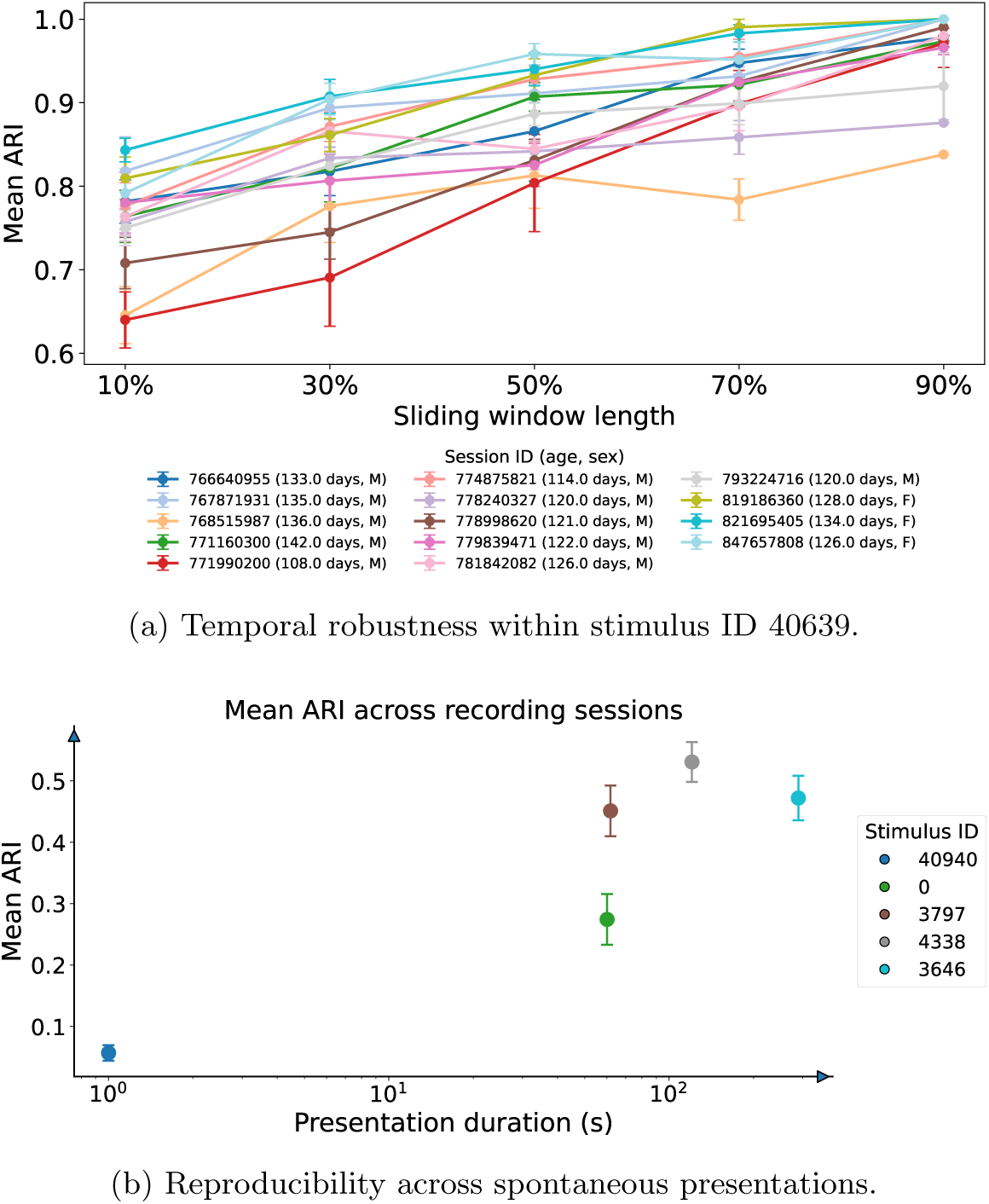
Temporal robustness and cross-presentation reproducibility of the vR-FCD communities in the Allen Neuropixels recordings. (a) Mean ARI between communities detected from subsampled windows of stimulus ID 40639 and those obtained from the complete recording, showing increasing agreement with window length across sessions. (b) Mean ARI (standard error) between the reference partition from stimulus ID 40639 and partitions obtained from the remaining spontaneous presentations, plotted against presentation duration. Longer recordings generally provide greater agreement, whereas the shortest presentation shows poor recovery.

### 5.2 Consistency of functional communities across spontaneous activity epochs

We next examine whether the community structure identified using the vR-FCD framework is reproducible across distinct spontaneous activity epochs within the same recording session. Using the partition obtained from stimulus ID 40639 as a reference (as this is the longest and most reliable block), we compare it against partitions computed from the remaining spontaneous presentations (Fig. 14 (b)). The mean ARI across all recording sessions are shown with error bars denoting the standard error. The results reveal a clear dependence on the duration of the spontaneous recording. The shortest presentation ID 40940 (1 s) exhibits almost no agreement with the reference partition. As the recording duration increases, the reproducibility improves substantially, reaching moderate to high agreement for presentations lasting approximately 60-121 s. The highest reproducibility is obtained for presentation ID 4338 (≈121 s), while the other long-duration presentations (IDs 3646 and 3797) also exhibit consistently high agreement. Although there is an overall positive relationship between recording duration and ARI, the dependence is not strictly monotonic. These results suggest that there exists a persistent intrinsic organisation of neuron communities across spontaneous activity epochs. The higher agreement for longer presentations suggests that a sufficient amount of data is required to recover the underlying community structure reliably, whereas low ARI for short epochs should not be interpreted as evidence of reorganisation of the neurons across the communities. Although the same units are tracked within each recording session, finite sampling, changes in spontaneous firing patterns, and possibly electrode drift or variation in spike sorting can all reduce cross-epoch agreement.

### 5.3 Relationship between functional communities detected and anatomical distribution of the neurons

Finally, we assess how the functional communities identified by our vR-FCD framework relate to the anatomical brain regions recorded in the Allen dataset, see Fig. 15. The detected communities are distributed across multiple cortical and subcortical regions, but clear regional biases are also visible. In several sessions, neurons in the visual cortical areas are predominantly associated with one functional community, whereas the thalamic regions are enriched for another, in spite of some mixing within individual regions. The hippocampal formation also shows regional organisation, and in some sessions is dominated by the same community enriched in visual cortex. This relationship is however not consistent across all recordings, as illustrated in Fig. S15 in the Appendix. The midbrain consists of mixtures of the detected communities. Thus the inferred functional organisation does not reproduce the anatomical distribution directly, but instead shows region-specific enrichment. This supports a partial correspondence between functional and anatomical organisation of the neurons. For more details on the anatomical distribution of the Allen data, see Appendix G.

**Figure 15:**
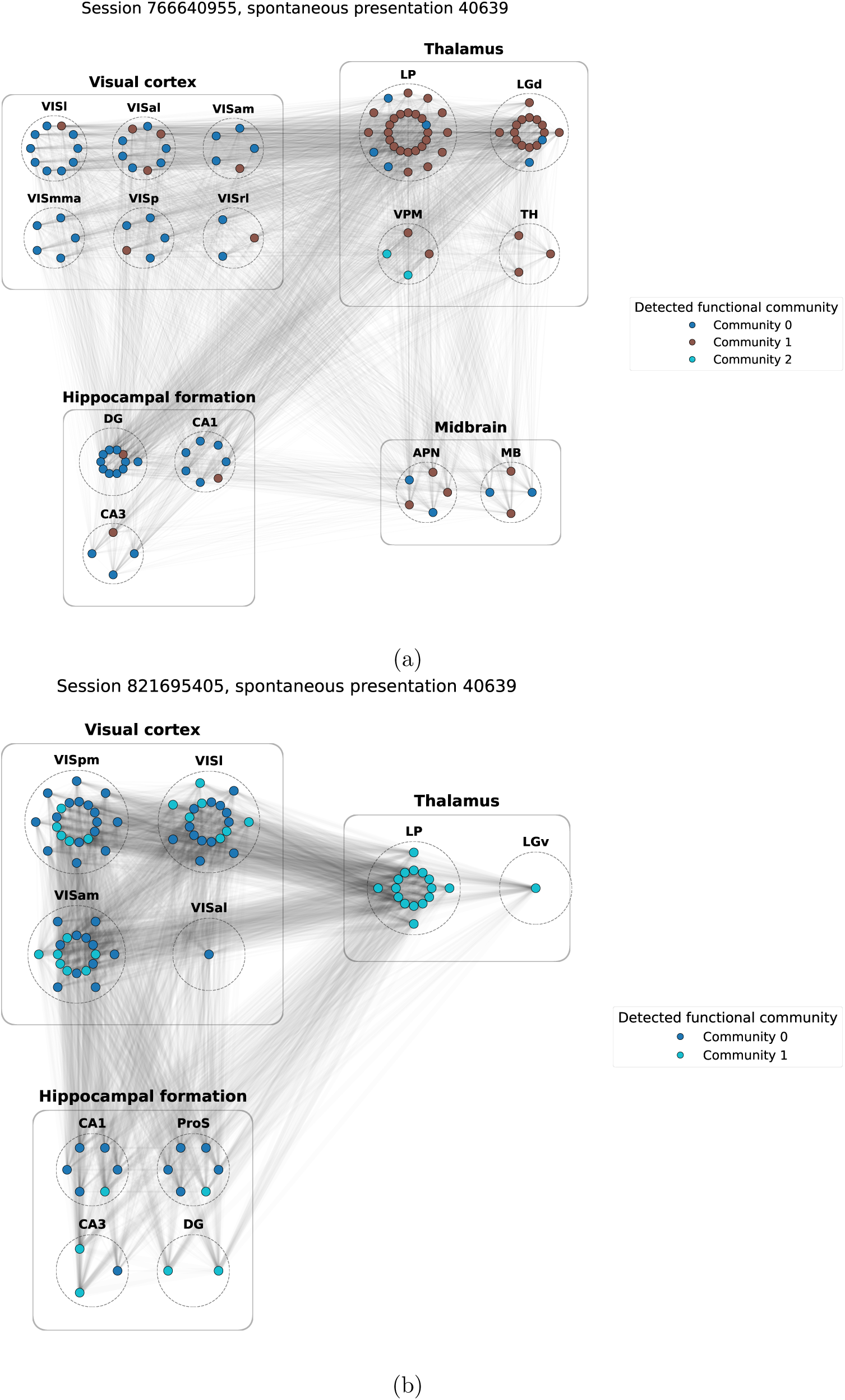
Functional neural assemblies detected using the vR-FCD framework from spontaneous data (ID: 40639) in two Allen sessions: (a) 766640955 and (b) 821695405. Nodes represent neurons coloured by detected community, dashed boundaries denote anatomical brain regions including the visual cortex, thalamus, the hippocampus, midbrain, etc., and edge weights are scaled by functional similarity. The recovered communities span multiple anatomical regions while exhibiting region-specific enrichment. This suggests distributed functional organisation with partial correspondence to anatomical structure.

## 6 Discussion and conclusion

In this work we introduce a framework for identifying functional communities directly from neural spike-activity data by combining spike train similarity (via van Rossum distance) with network-based community detection using the Louvain method which is called **vR-FCD** (van Rossum-Functional Community Detection). We transformed the spike-activity time series data into a functional similarity network whose community structures revealed groups of neurons exhibiting coordinated dynamics. Broadly speaking, this work provided a bridge between spike-train analysis and network science, offering a principled and computational approach for uncovering latent organisation in neural populations. Our approach provides a systemic way to study how mesoscale communities of neurons emerge, reorganise, and persist across different temporal scales and experimental conditions. Although the similarity network in our approach lies in a latent space representing a functional network, it is static unlike the temporal functional network in Vaiana *et al* [30].

Overall, the vR-FCD framework performs well when neuron communities are sufficiently distinct. In synthetic data, recovery improves with larger phase separation, lower jitter, and longer recordings. The analyses of the SBM network of the LIF neurons show similarly strong recovery when structural connectivity and external drive produce clearly differentiated cluster dynamics. The robustness analyses further show that the main conclusions are not sensitive to the choice of similarity transformation. Additional experiments on the synthetic data includes the interaction between ISI and phase separation (Appendix C.3). For fixed phase separation, increasing ISI mainly reduces recovery for short recordings and higher jitter because spikes are available for estimating pairwise similarities, whereas longer recordings largely remove this dependence. In the high-noise regime, recovery is instead governed primarily by the phase geometry. Across both the synthetic data and LIF simulations, the clipping, exponential, and min-max transformations reproduce the same qualitative behaviour with only modest quantitative differences see Appendix C and F. For the Allen Neuropixels data, where no ground truth is available, the inferred communities remain largely consistent across temporal subsamples and show partial correspondence with anatomical organisation.

A broader application of our methodology is to identify and analyse functional neural assemblies directly from spontaneous spike activity. Such assemblies are relevant to questions of sensory processing, information integration, memory, and coordinated population dynamics, where the organisation of neuron groups may change across brain states or behavioural conditions [31]. The framework therefore provides a way to track how these neuron assemblies change over time. One specific application is epilepsy, where seizures are widely regarded as emergent phenomena involving abnormal synchronisation across neural populations [32]. In this context, vR-FCD could be used to examine how functional communities reorganise before, during, and after seizure events. This could potentially reveal network-level changes that are not apparent from individual neurons alone.

The framework has several limitations that also point to clear directions of extension. Reliable recovery requires sufficiently long spike-train recordings. Short observations, high noise or weak dynamical separation between communities can substantially reduce performance. This is particularly relevant for experimental data, where functional assemblies may evolve over time rather than remain stationary within a recording window. Although our sliding-window analysis provides evidence of temporal robustness, a more appropriate representation of changing assemblies would be a temporal or multiplex network in which the same neurons are linked across successive windows, allowing community evolution to be tracked directly. A second limitation is that functional communities need not coincide with anatomical organisation. The partial correspondence observed in the Allen data is therefore informative, but temporal reproducibility alone cannot establish that the inferred communities represent a unique biological ground truth. In addition vR-FCD currently relies on pairwise, symmetric van Rossum similarity, so directional influence, causal interactions, and higher-order coordination among groups of neurons are not captured. Incorporating directed measures, alternating spike-train similarity metrics like the *Victor-Purpura distance* [33, 34, 35], and higher-order network representations could therefore provide a richer description of neural interactions. The use of modularity-based community detection also introduces resolution limitations, motivating consensus-based approaches that could reveal multi scale organisation and quantify uncertainty in community assignments. Finally, constructing the full *N × N* distance matrix becomes increasingly expensive for large populations and long recordings. GPU acceleration, dimensionality reduction, and more scalable network methods would therefore be important for extending vR-FCD to larger scale electrophysiological datasets.

Our framework provides a practical framework for uncovering the mesoscale functional organisation directly from spike trains, with robust recovery when sufficient data and clear dynamical separation are present. This offers a foundation for extending community based analysis to increasingly large and time dependent neural recordings.

## Code availability

The `Python` codes that support the findings of this study are openly available from the GitHub link `https: //github.com/indrag49/vanRossum-Network`. The python package `vrfcd` developed for this study is open source and available on GitHub at https://github.com/indrag49/vrfcd. The package is also available to download from the Python Package Index (PyPI) via `pip install vrfcd`.

## Acknowledgements

This work was supported by a grant (#22/FFP-P/11547) managed by Research Ireland. The authors acknowledge the Research IT HPC Service at University College Dublin for providing computational facilities and support that contributed to the research results reported in this paper. IG thanks Yi Wang from Massey University for useful discussion.

## AI use

ChatGPT-5.5 Plus was used to assist with debugging, edge case handling, plotting aesthetics, and generating initial code structures with complete human oversight.

## A Quantitative measures integrated in the vR-FCD framework

Here we go into more detail about the methodology by listing the quantitative measures used to compare spike trains and identify functional communities: the van Rossum distance, the construction of a weighted functional similarity network, and Louvain community detection that integrates into the vR-FCD framework. We also discuss ARI as an established statistical measure to assess accuracy.

### A.1 van Rossum distance

Given a population of *N* neurons, let 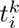 denote the time of the *k*-th spike for neuron *i*. Let the spike train be represented by the collection of spike times 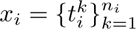 for *i* = 1*,. .., N* where *n_i_* is the total number of spikes in neuron *i* (Fig. S1 (a)). The van Rossum distance compares spike trains after filtering them with an exponential kernel

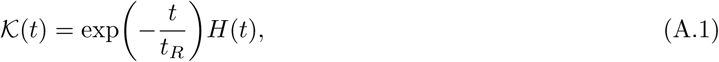

where *H*(*t*) is the Heaviside function and *t_R_* defines the characteristic temporal scale of the exponential kernel (Fig. S1 (b)). The parameter *t_R_* controls the tightness of synchrony that is captured by the distance measure: small values emphasise only very closely aligned spikes, whereas larger values allow similarities over broader temporal windows. A common choice of *t_R_* is to select it relative to the characteristic average ISI [35]. In practice, choosing *t_R_* to be approximately one order of magnitude smaller than the mean ISI often provides a good balance. The van Rossum distance is

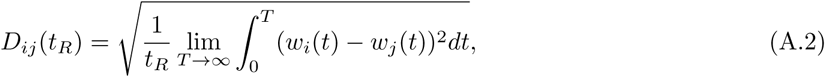

**Figure S1:**
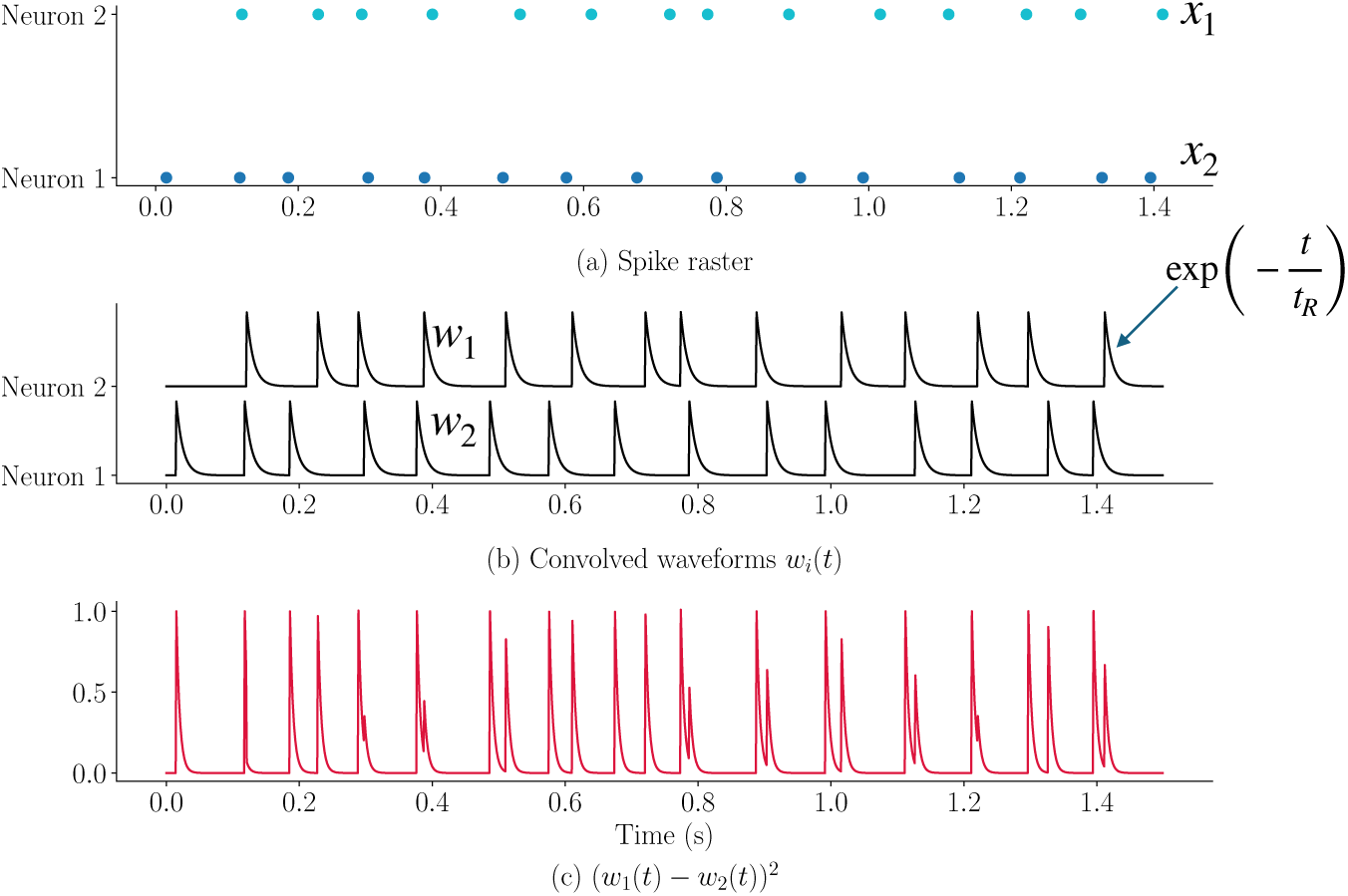
Constructing the van Rossum distance from spike trains. Two example spike trains. (a) are converted with an exponential kernel to produce the waveforms *w*_1_(*t*) and *w*_2_(*t*) (b). Their squared difference over time (c) provides a measure of spike-train dissimilarity at the temporal scale *t_R_*.

where *w_i_*(*t*) is the convolution

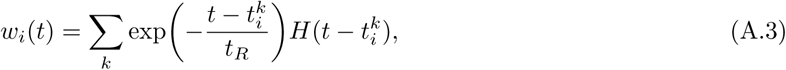

(Fig. S1 (c)). Following van Rossum [17], we normalise the distance by the square root of the average number of spikes in the two spike trains,

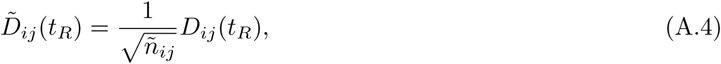

Where 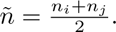 When two neurons *i* and *j* are fully synchronised, i.e. their spike times will be identical and *D_ij_* = 0. A value *D̃_ij_ >* 0 represents the degree of mismatch between the two spike trains, i.e. asynchrony. Although the normalisation is intended to place *D̃_ij_* approximately in the interval [0, 1], it is not a strict upper bound. Since, the value *D̃_ij_* is normalised by the average number of spikes between the two spike trains, rather than by a strict theoretical maximum, two neurons with very different spike rates can produce disproportionately large waveform differences after convolution, causing *D̃_ij_* to slightly exceed 1.

### A.2 Computation of the functional similarity matrix

We convert the van Rossum distance matrix into a weighted similarity network using the *min-max* normalisation

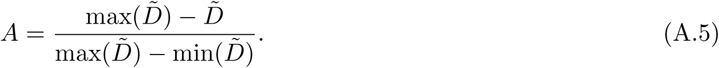

which maps the distances to the interval [0, 1], with larger values corresponding to greater spike-train similarity. The resulting functional network is fully connected and weighted with nodes representing the neurons and the edges quantifying functional similarity. We remove the self loops by setting *A_ii_* = 0. Besides the min-max transformation, we have tested other transformation functions on the distance matrix *D̃* also:

i. After running several experiments with different synthetic spike rasters, we note that for two spikes which are highly asynchronous or dissimilar, the distance can be *>* 1 however it never crosses 1.1. Thus, it is safe to consider a threshold value of 1 and clip any *D̃_ij_* value to 1 if it is just slightly higher than 1. After constructing the van Rossum distance matrix, *A* is given by

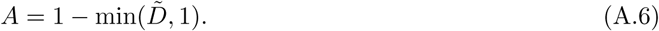
ii. The clipping over 1 can be avoided altogether by applying the threshold function

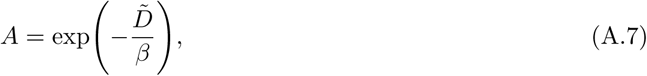

where *β >* 0 is a scaling parameter which needs to be fixed based on simulation experiments. Note that *A*(*t_R_*)_max_ = 1, as for any *x* ≥ 0, 0 *<* exp(*x*) 1. The thresholding function is similar to the *heat kernel* function used by Belkin and Niyogi [36] for weighting the edges in a network.

Across synthetic data, LIF network simulations, and Allen recordings, the min-max transformation provided the most consistent overall performance. We therefore use it as the primary transformation in our analyses for the following reasons:

i. Even though all three schemes work similarly for the synthetic data, and the data simulated using LIF neurons on a SBM network, it is just the min-max scheme which seems to reasonably work for the Neuropixels data after several testings.
ii. The min-max normalization adapts to the data at hand, by using actual distribution of the distances in *D̃*. There also exists automatic rescaling for different duration *T* and number of neurons *N*.
iii. The clipping of the values to 1 in (A.6) is arbitrary and does not provide a general setup, and eventually loses information for values of *D̃ >* 1.
iv. For the exponential function (A.7), there exists another parameter *β* which needs additional tuning and adds and extra label of complexity. Also, many values transforms to 0 and the distribution of the values become highly skewed.
v. Also, in terms of applying the Louvain algorithm, the min-max normalization produces well-spread weights that leads to better community partitions, unlike the other two transformations.

### A.3 Louvain algorithm for community detection

Louvain algorithm is a community detection algorithm that depends on optimising the network *modularity* [37, 38]. For a weighted network, the modularity is given by

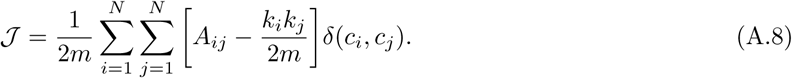

Here *A_ij_* represents an element of the adjacency matrix *A*, i.e. *k_i_, k_j_* represent the sum of the weights of all links connected to nodes *i* and *j* respectively: 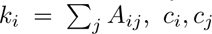 are the communities to which node *i* and *j* belong respectively, and *m* is the sum of all the link weights of the weighted network, i.e. 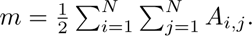. Also, *δ*(.,.) represents the Kronecker-delta function.

The algorithm runs in two phases that are iterated until no further increase in network modularity is possible. It starts by assigning a distinct community to each of the *N* nodes of the weighted network (Fig S2 (a)). For each node *i* the change in modularity ΔJ is computed for moving it to a neighbouring community *c_j_* from its own community *c_i_*. Nodes are then reassigned to communities that offer the highest and positive ΔJ. This process iterates across the network until *Q* stabilises at a local maximum (Fig S2 (b)).

**Figure S2:**
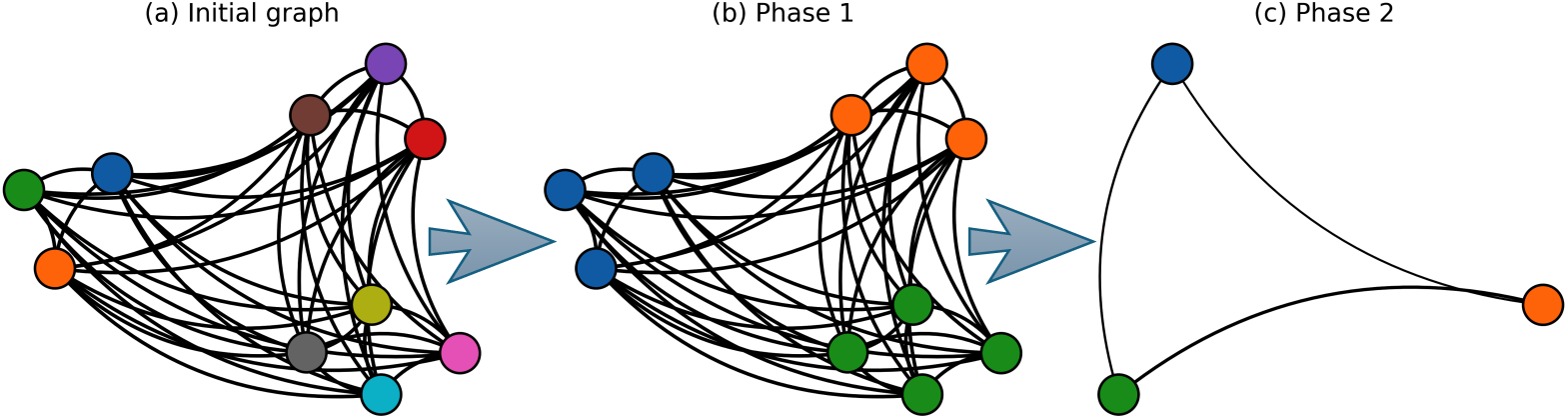
A sample visualization of the two phases of the Louvain algorithm. It starts by an initial assignment of a distinct community to each node. In the first phase the nodes are iteratively reasigned to communities to maximize the modularity J, giving rise to local clusters. In the second phase the communities are aggregated into single nodes, producing a reduced graph on which the process is iterated until there is no gain in modularity. This is the final community structure.

Now, in the second phase, a new network is built, where individual nodes that constitute a single community in the first phase are aggregated into a single node. The weight of the link between two such nodes is the sum of the weights of the links between the corresponding communities in the first phase (Fig. S2 (c)). This process consisting of the two phases is repeated on the new network until there are no more changes and the optimum modularity has been obtained. We apply the Louvain algorithm on the functional similarity matrix to detect communities. The Louvain algorithm provides an efficient method for detecting modular structure in large weighted networks.

### A.4 Accuracy measure (ARI)

The *adjusted Rand index* (ARI) is used to validate whether the communities are properly detected, given the ground truth is known beforehand. The concept of ARI builds on the *Rand index* (RI) [22] named after William M. Rand, which gives a score on how similar two clustered datasets are. Wagner and Wagner [39] defined the comparison between two clusterings in terms of the number of pairs of elements that are assigned to the same or different clusters in both partitions. The Rand index ranges from 0 to 1, taking the value of 1 when the two clusterings are identical and 0 when all pairwise assignments between the clusterings are completely mismatched. However, RI exhibits an issue of overestimating the quality of the clusters. i.e. counting agreements that occur purely by chance. Hubert and Arabie [40] proposed the ARI where the raw RI is adjusted by subtracting the expected similarity from random chance and normalising by the maximum possible similarity. For a perfect match between clusters, the ARI takes a value of 1. Note that it can be negative when the results of the clustering obtained are worse than what is expected by random choice. It takes a value of 0 for random labelling independently of the number of clusters.

## B Simulations using Python

We have used the `convolve()` function from the signal processing package `scipy.signal` for convolving our discrete spike trains as a step in determining the van Rossum distance. For the network theory segment of our work we have used the popular `NetworkX` package that lets us create, study and analyse complex networks in `Python`. For implementing the Louvain algorithm we can use the `louvain communities()` function provided by `networkx.algorithms.community` that is a collection of functions for computing and measuring community structures in complex networks. The parameters that the `louvain communities()` requires are the `weight` of the edges which we input from the transformed adjacency matrix, the `resolution` which is by default kept at 1 (the algorithm favour larger communities when *<* 1, and favours smaller communities when *>* 1), the `threshold` which by default is 10*^−^*^7^ but can be provided with even smaller values which by far did not make any difference, the `max level` which takes the maximum number of steps of algorithm and is kept `None` by default (meaning the stopping condition of the algorithm is determined by the `threshold`), and the random `seed`. Finally to implement ARI on the predicted clusters we utilise the `scikit-learn` package which is a machine learning toolkit for `Python`. It is imported as `sklearn` and provides different score functions and performance metrics through its `sklearn.metrics` suite. Specifically we use the `adjusted rand score()` that lets us compute ARI to validate the clustering procedure. Also, to improve the performance of the *N XN* computations and to make large parameter sweeps feasible, we implemented parallel processing using the `joblib` library through the `Parallel()` and `delayed()` functions to distribute independent simulations and distance computations across multiple CPU cores.

## C Application of the vR-FCD framework to the synthetic data

### C.1 Synthetic data preparation

A neuron’s *k*−th spike from block *b* happens at time *t^k^*:

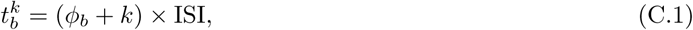

where *ϕ* moves the entire spike time forward in time and controls the relative phase between two sequential blocks in the synthetic data. This is how we introduce relative phase differences between two blocks. This parameter *ϕ* is dimensionless, and is specified as a fraction of the ISI. Next, we use the jitter parameter *σ* to control the amount of temporal noise added to the otherwise periodic spike train (C.1). This parameter *σ*, like *ϕ* is also dimensionless and is specified as a fraction of the ISI. We define the maximum jitter in time bins as

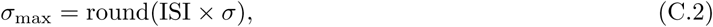

where the *round* function is defined by round(*x*) = ⌊*x* + 0.5⌋. Every spike in a spike train is random integer uniformly sampled from [−*σ*_max_*, σ*_max_]. An actual spike time can be updated from (C.1) to

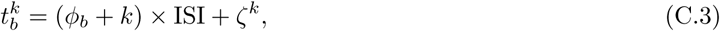

where *ζ^k^* ∼ *U* [−*σ*_max_*, σ*_max_].

### C.2 Effect of phase separation *ϕ* and jitter *σ* on the accuracy value ARI for the clipping and exponential kernels

Next, we show the plot of ARI as function of *ϕ* with varying *σ* for the clipping transformation (Fig. S3). Highest recovery occurs when the block phases have high phase differences between the blocks. Recovery decreases where the phases between the blocks overlap. With high *σ* the range of perfect recovery decreases and longer recordings also stabilise recovery. The behaviour is similar for the clipping transformations.

**Figure S3:**
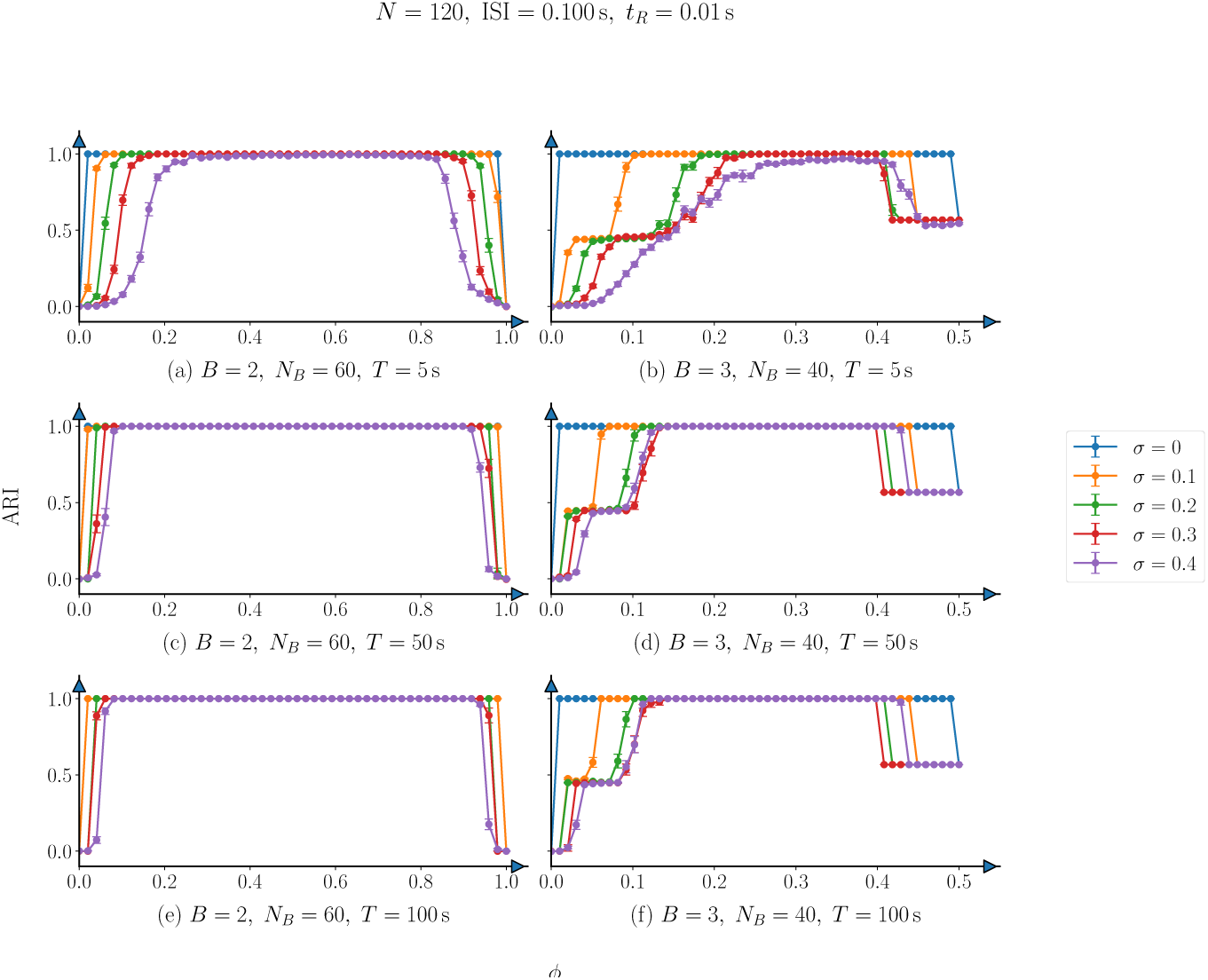
ARI as a function of total number of phase separation *ϕ* for the clipped transformation. Here ISI = 0.1 s, and *t_R_* = 0.01 s. Different curves correspond to *σ ∈* [0, 0.4]. Recovery is the highest when the block phases are well separated and decreases near the boundaries where the phases overlap. Larger *σ* reduces the range of perfect recovery. For *B* = 3, the cyclic phase geometry causes ARI to decrease again when the first and third blocks become similar. Longer recordings stabilise recovery.

### C.3 Effect of inter spike interval (ISI) and phase separation *ϕ* on the accuracy value ARI

At fixed high jitter *σ* = 0.4 and *T* = 100 s, community recovery depends strongly on the phase geometry (Fig. S4). For *B* = 2, well separated phases yield near-perfect recovery across the ISI range, whereas *ϕ* = 0, 1 give *ARI* ≈ 0 because the two-blocks are phase aligned. For *B* = 3, moderate phase separations are recovered well, while *ϕ* = 0 fails and *ϕ* = 0.5 gives partial recovery because the first and the third blocks overlap. Across the three similarity transformations, the min-max and exponential cases remain comparatively stable with ISI, whereas the clipping transformation shows a sharper deterioration at larger ISI.

**Figure S4:**
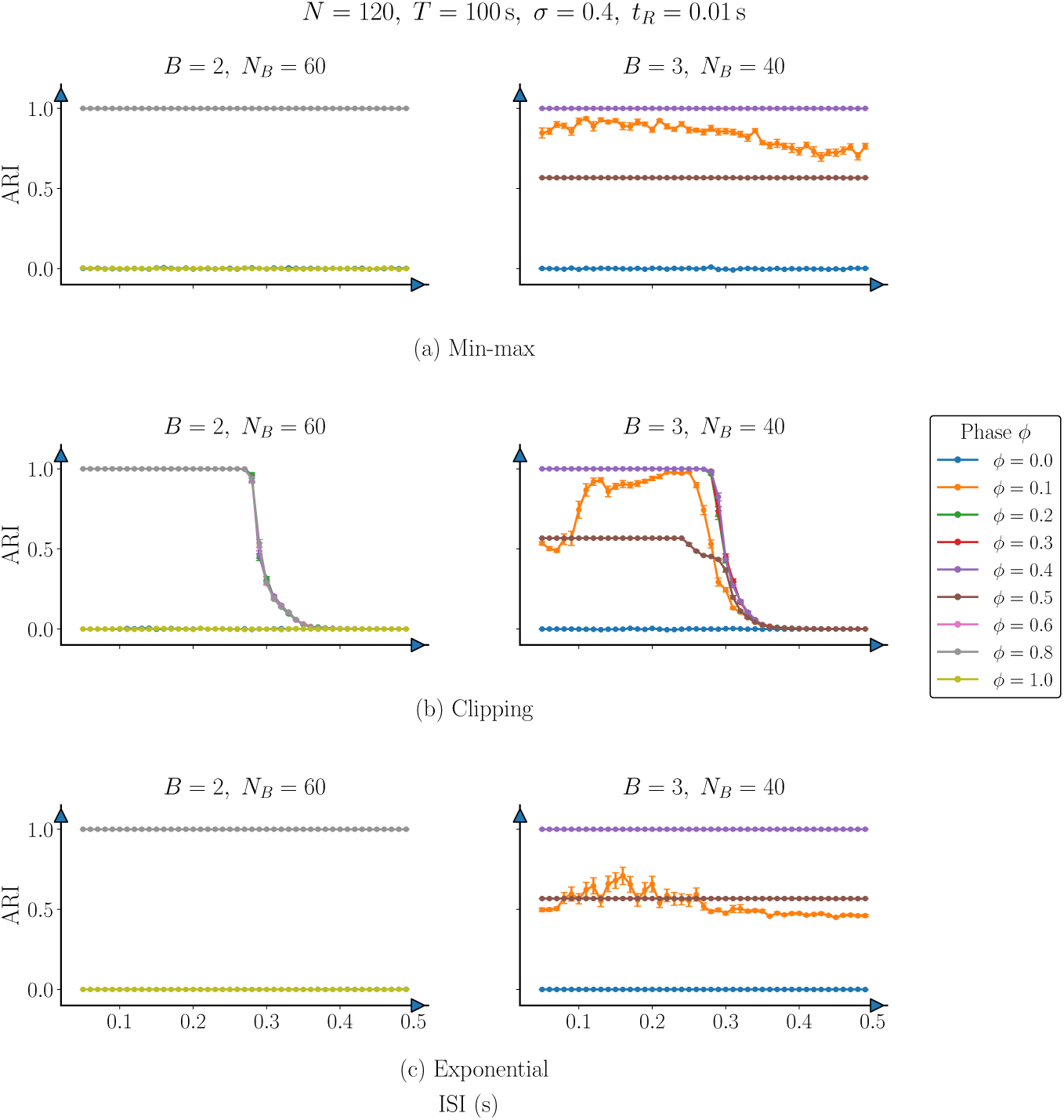
ARI as a function of ISI for a fixed high jitter value *σ* = 0.4, for (a) min-max, (b) clipping, and (c) exponential transformations. Here *T* = 100 s, *N* = 120, and *t_R_* = 0.01 s. Curves correspond to different phase separations *ϕ*. For *B* = 2 well-separated phases give near-perfect recovery whereas *ϕ* = 0, 1 produce ARI 0 due to phase overlap. For *B* = 3, moderate phase separations gives high recovery while *ϕ* = 0 fails and *ϕ* = 0.5 gives partial recovery because the first and third blocks overlap. The min-max and exponential transformation remain stable across ISIs, whereas clipping shows a shard deterioration at larger ISI.

### C.4 Effect of network size *N* and varying splits on the accuracy ARI

In this section, we examine how vR-FCD performance varies with network size *N* and community balance under the clipping and exponential transformations (Fig. S5). For *B* = 2, recovery rapidly approaches *ARI* = 1 as *N* increases, whereas *B* = case is more sensitive to community splits. Balanced partitions generally achieve better recovery. Both transformations reproduce the same qualitative dependence on *N* and community splits.

**Figure S5:**
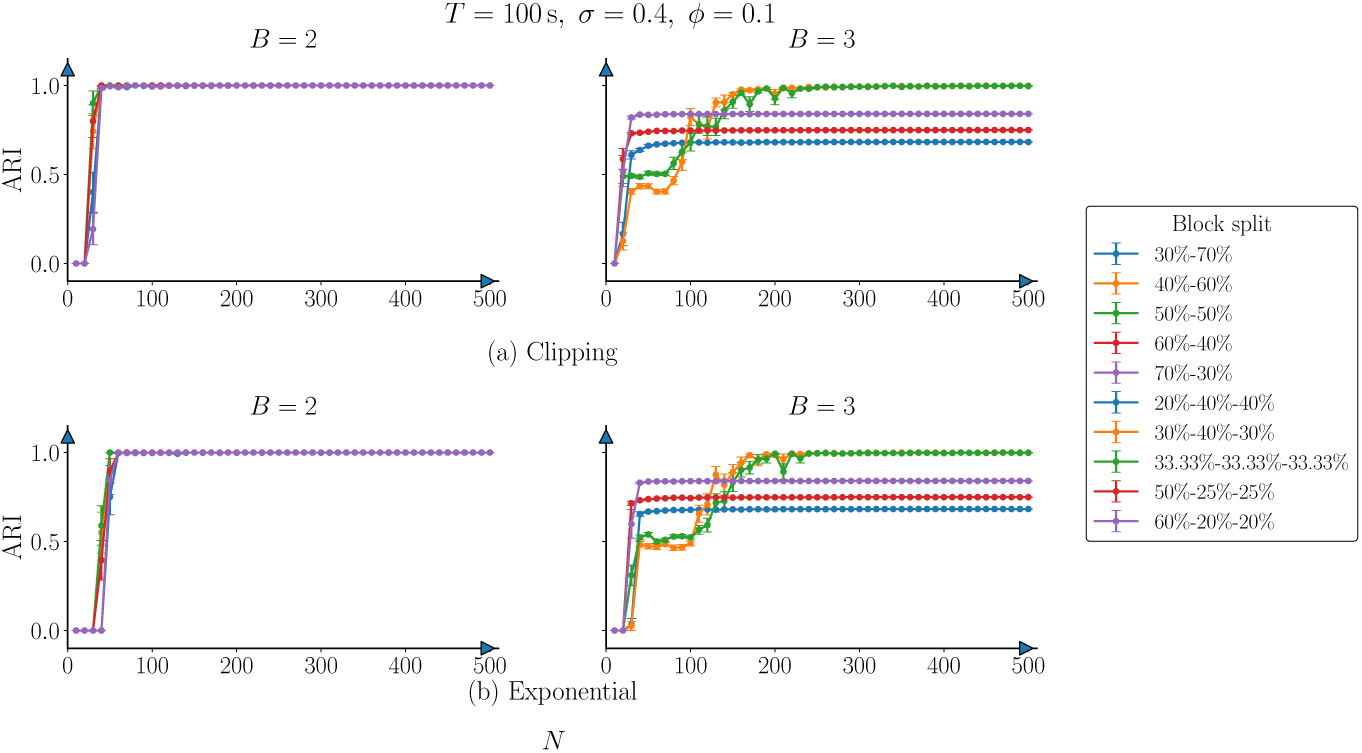
ARI as a function of total number of neurons *N* for different size splits under the transfor mati on (a) *A* = 1 − (*D̃*, 1) with clipping at **1** and (b) the exponential transformation 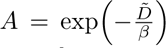 with β = 0.1. Here T = 100 s, ISI = 0.1 s, σ = 0.4, and ϕ = 0.1. For B = 2, both transformations rapidly achive near-perfect recovery as *N* increases. For *B* = 3, recovery depends more strongly on the community splits. Balanced splits improve with *N* and can reach ARI 1, whereas strongly imbalanced ones exhibit low ARI values.

### C.5 Effect of the van Rossum time-scale parameter *t_R_* and noise levels *σ* on the accuracy ARI

We next examine how community recovery depends on the van Rossum time scale *t_R_*at *T* = 100 s for different *σ* (Fig. S6). For the clipping transformation, recovery can change abruptly as *t_R_* increases, particularly at higher *σ*, whereas the exponential transformation produces a more gradual decline. The *B* = 3 case is consistently more sensitive than *B* = 2, reflecting smaller temporal separation between neighbouring blocks. Both transformations confirm that the choice of *t_R_* becomes important with increase in noise and community overlap.

**Figure S6:**
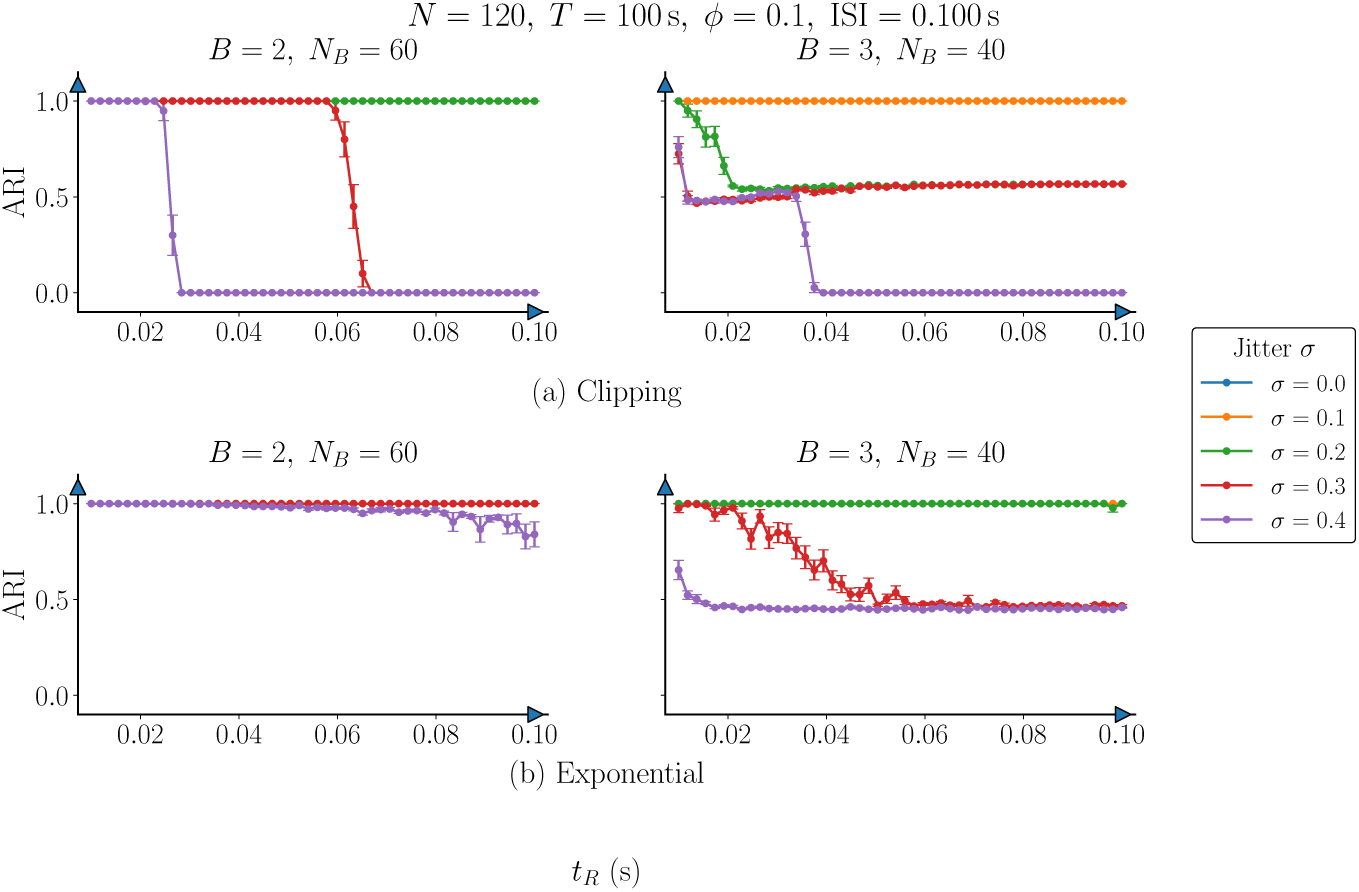
ARI as a function of the van Rossum time-scale parameter *t_R_* for different *σ* values under the transformati on *A* = 1 − (*D̃*, 1) with clipping at 1 (panel (a)) and the exponential transformation 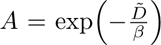 with β = 0.1 (panel (b)). Here N = 120, T = 100 s, ϕ = 0.1 s. The clipping transformation exhibits sharp transitions in recovery as *t_R_*increases, particularly at higher *σ*, whereas for the exponential transformation the changes are more gradual. The *B* = 3 case is more sensitive to both *t_R_* and *σ*, reflecting the smaller separations between neighbouring communities.

## D A network of Leaky Integrate and Fire (LIF) neurons

In § 3 we apply the vR-FCD framework to spiking data from a simulation of Leaky Integrate-and-Fire (LIF) neurons. Here, we describe the LIF network and how this data is generated. For a neuron *j* in a particular network topology, voltage *V_j_* is given by

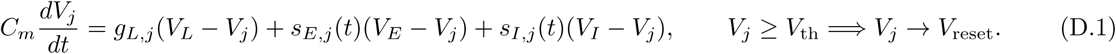

Neurons can be either excitatory (*E*) or inhibitory (*I*) depending on the type of neurotransmitter they release at their synapses [41]. In (D.1) *C_m_*is the membrane capacitance which determines how fast the voltage changes, *V_j_* is the membrane potential of a neuron, *V_L_*is the resting potential, *V_E_* is the excitatory reversal potential, *V_I_* is the inhibitory reversal potential, *g_L_* is the leak conductance (excitatory neurons have a different value of leak conductance as compared to the inhibitory neurons implying a functional asymmetry in the types of neurons), *s_E,j_* is the synaptic excitatory conductance, and *s_I,j_* is the synaptic inhibitory conductance. In between spikes, the synaptic conductances decay exponentially. A spike is detected when the membrane potential of a neuron *j* at time *t* denoted by *V_j_*(*t*) crosses a threshold *V*_th_. It is then reset to a value *V*_reset_.

The update in the excitatory synaptic conductance is

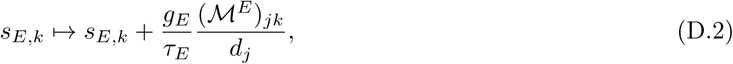

Each neuron *j* receives an external drive given by the Poisson point process

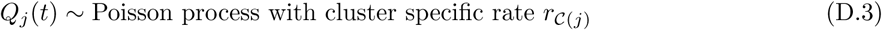

The firing rate *r_C_*(*j*) is cluster dependent with unit Hz. The drive is input to excitatory conductances only.

So we can write down the synaptic dynamics as

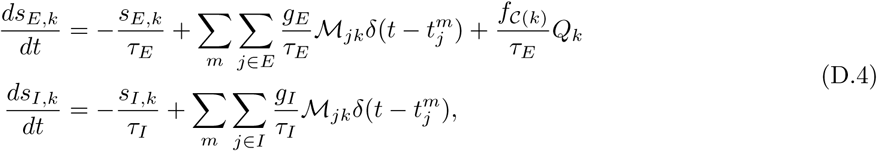

where the average cluster specific input is *f_C_*_(*j*)_ × *r_C_*_(*j*)_. Here *t^m^* is the time when the neuron *j* spikes the *m*^th^ time. Some of the fixed parameters considered in this paper are found in Tab. S1.

**Table S1:**
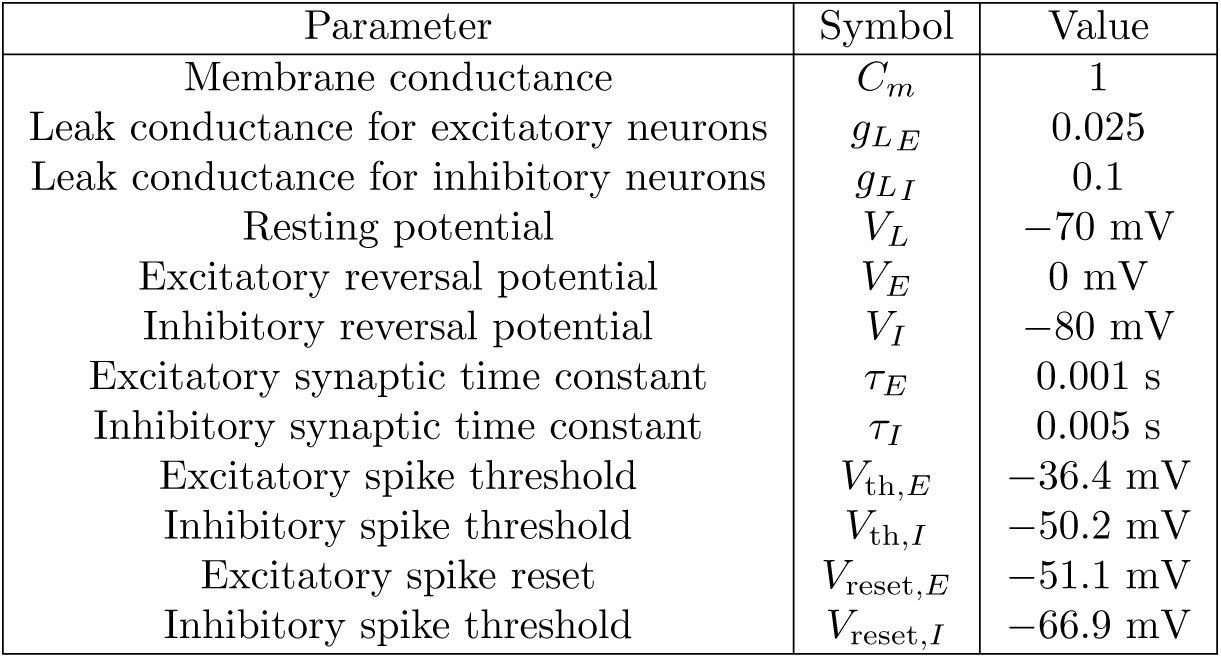
Fixed parameters for the conductance based Leaky Integrate and Fire (LIF) neurons. The excitatory neurons have slower, more integrative leaks associated to them whereas the inhibitory neurons have faster, more responsive leaks associated to them. The synaptic time constants imply fast excitation and slower inhibition, stabilising the network activity.

## E Algorithm outlining the application of vR-FCD to the the stochastic block model (SBM) network of Leaky Integrate and Fire (LIF) neurons

## F Application of the vR-FCD framework to the stochastic block model (SBM) network of Leaky Integrate and Fire (LIF) neurons

Algorithm 1 provides a detailed outline of application of vR-FCD to the the stochastic block model (SBM) network of Leaky Integrate and Fire (LIF) neurons and measuring accuracy value ARI. For two-dimensional parameter sweeps for the clipping and exponential transformations using two-cluster planted structure see Fig. S7 and for all three transformations using a three-cluster planted structure see Fig. S8. A different set of fixed parameters to portray similar two-dimensional parameter sweeps as was done in the last two figures are shown in Fig. S9 and Fig. S10. We also visualise the performance of vR-FCD as a function of the network size *N* for two-cluster planted structure using the clipping and the exponential transformations (Fig. S11), and for three-cluster planted structure using all three transformations (Fig. S12). We visualise the performance of vR-FCD as a function of the time-scale parameter *t_R_* for two-cluster planted structure using the clipping and the exponential transformations (Fig. S13), and for three-cluster planted structure using all three transformations (Fig. S14).

**Figure S7:**
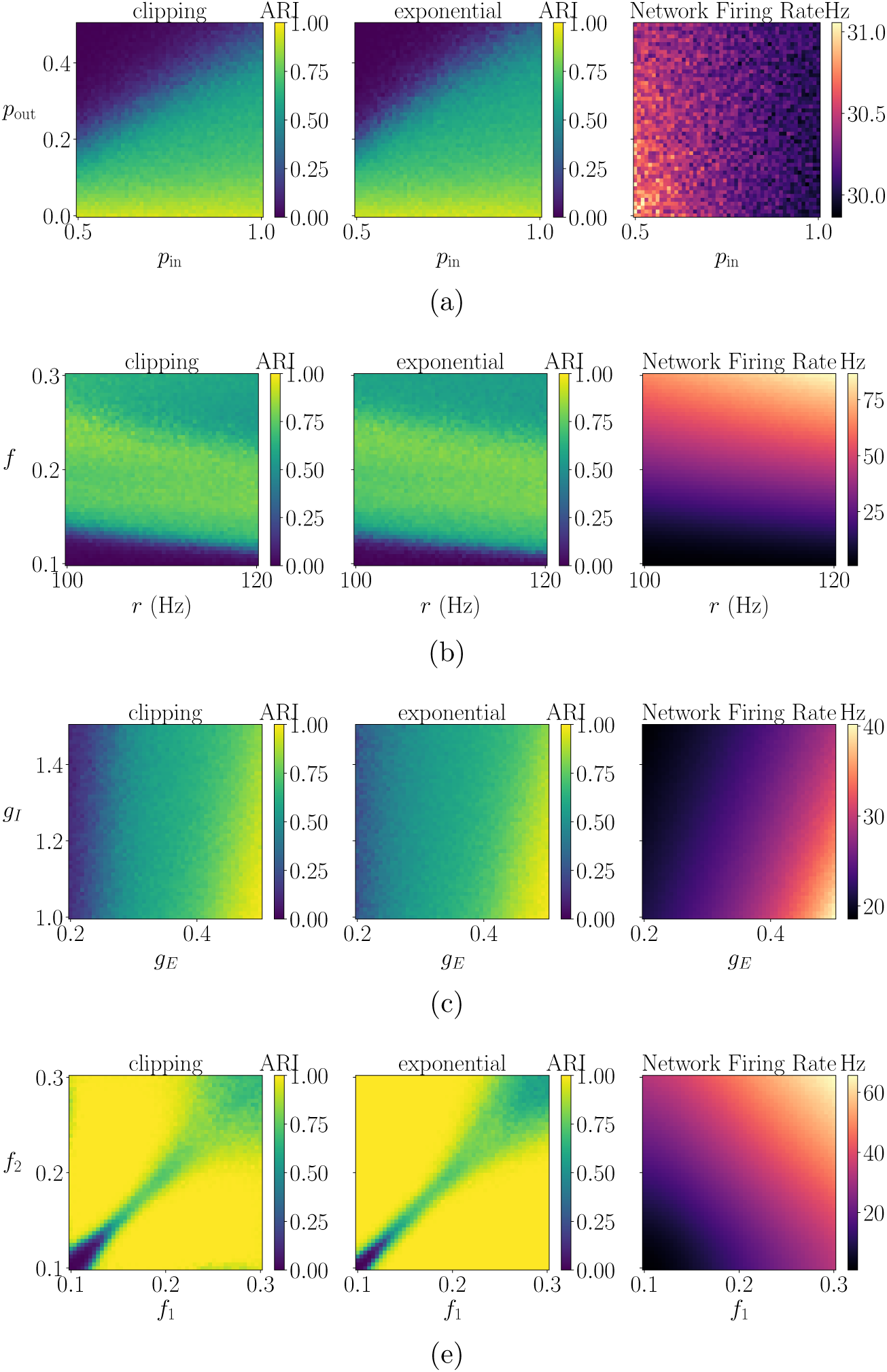
Two-dimensional parameter sweeps showing the mean ARI (left and the middle columns for the clipping and the exponential transformations respectively) and the firing rate (right column) for the two-cluster SBM. Here *t_R_* = 0.005 s. Other parameters are set as following Tab. S1 with *T* = 20 s, Δ*t* = 0.0001 s. Recovery is the highest when the two communities remain dynamically distinct, especially for high *p*_in_ and heterogeneous external drive. Recovery decreases when their dynamics become more homogeneous.The firing-rate plot indicates that recovery is not directly related to the changes in the average activity level.

**Figure S8:**
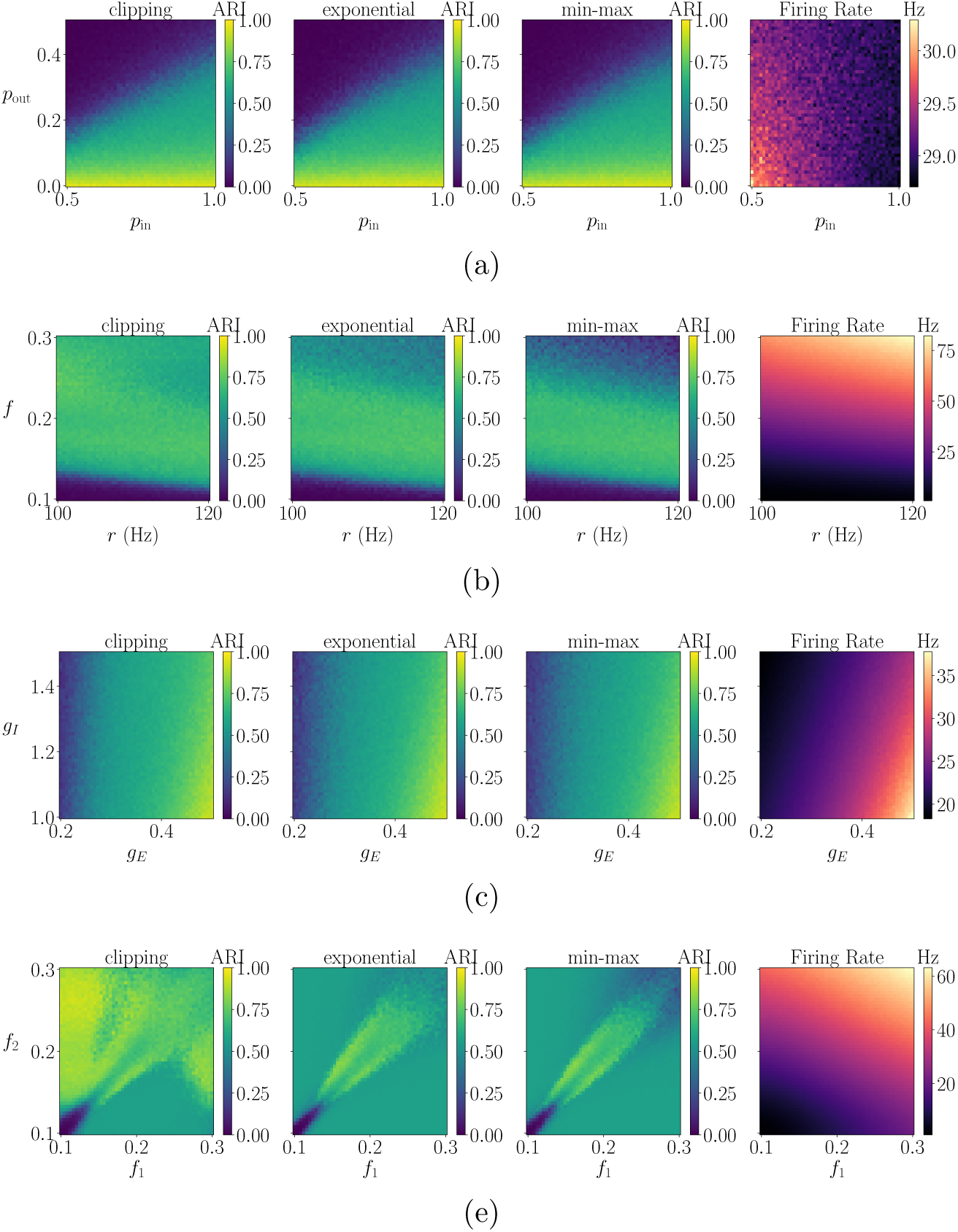
Two-dimensional parameter sweeps showing the mean ARI obtained using the clipping, exponential, and the min-max transformations (the first three columns respectively), together with the firing rate (right column) for the three-cluster SBM. Here *t_R_*= 0.005 s. High performance occurs in regimes where the three planted blocks remain dynamically distinguishable, especially for high *p*_in_ and sufficiently heterogeneous external drives. The clipping transformation is more sensitive to overlap between communities. The firing-rate plot indicates that changes in the recovery is not directly related to the changes in the average activity level.

**Figure S9:**
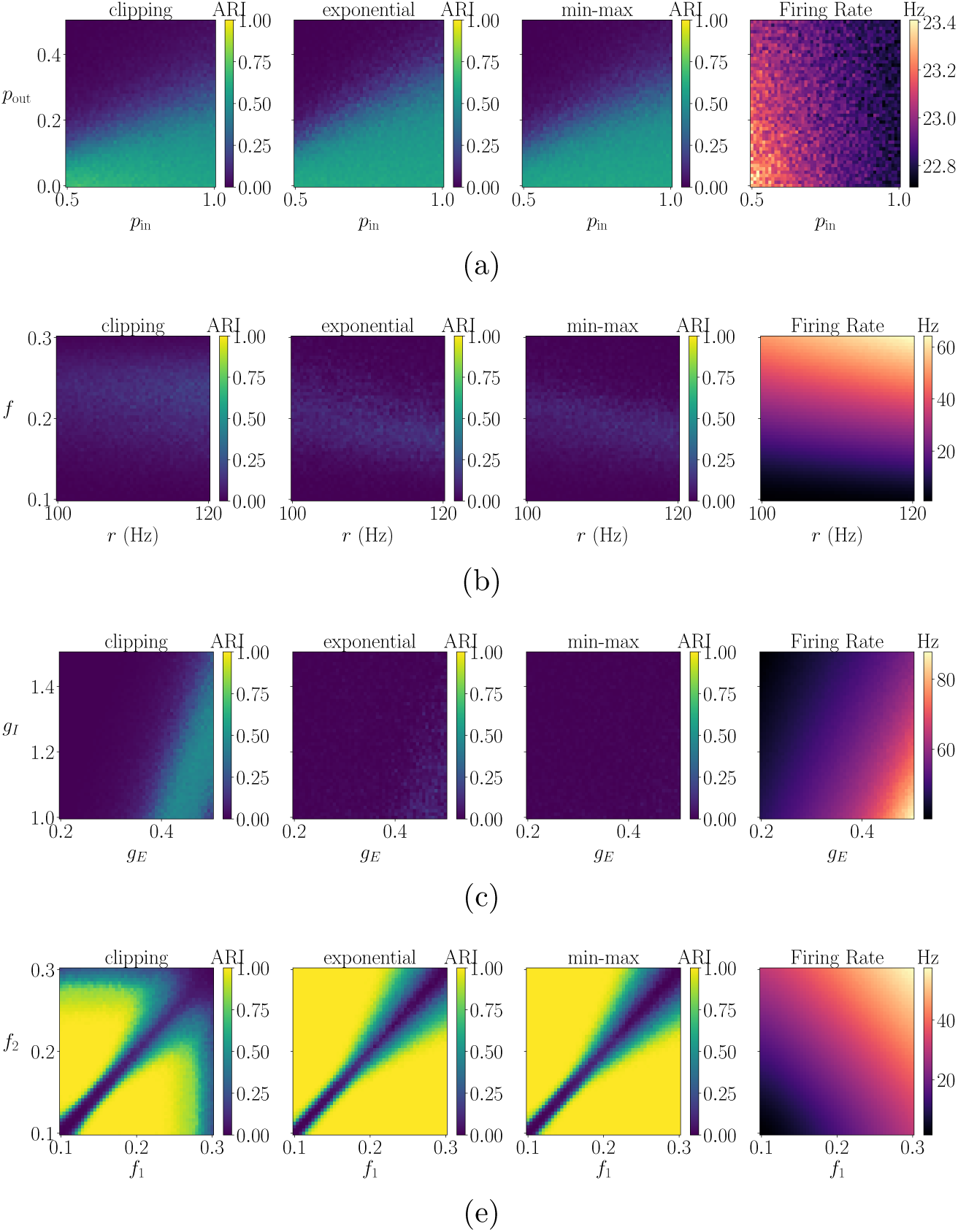
Two-dimensional parameter sweeps showing the mean ARI obtained using the clipping, exponential, and the min-max transformations (the first three columns respectively), together with the firing rate (right column) for the two-cluster SBM. Here *t_R_* = 0.005 s. Other parameters are set as following Tab. S1 with *T* = 20 s, Δ*t* = 0.0001 s. Recovery is generally reduced by stronger *p*_out_ and inhibition, which make the two planted communities less dynamically distinguishable. However, differences in *f*_1_ and *f*_2_ can restore separation, producing higher ARI away from *f*_1_ *f*_2_. The firing-rate plot indicates that changes in the recovery is not directly related to the changes in the average activity level.

**Figure S10:**
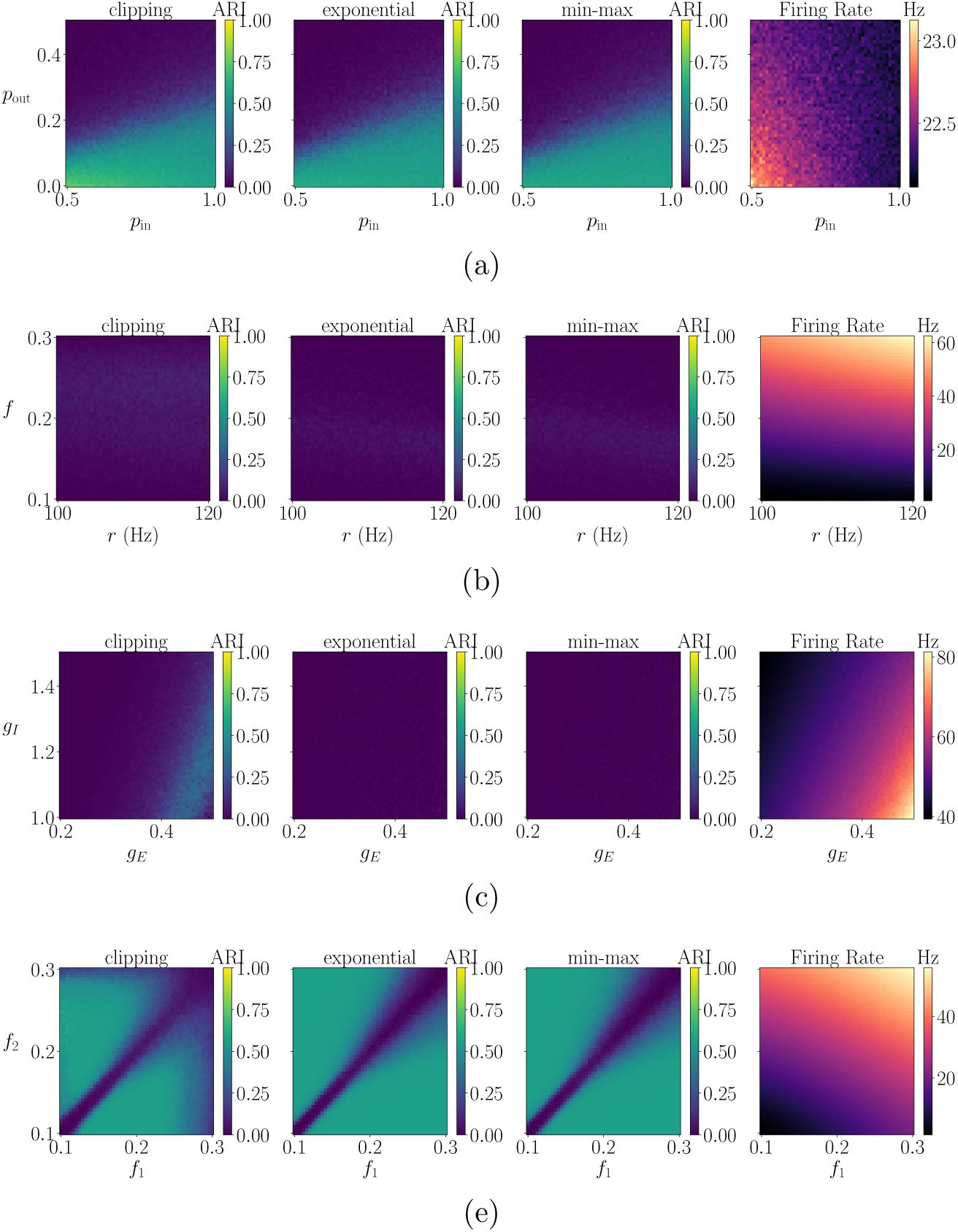
Two-dimensional parameter sweeps showing the mean ARI obtained using the clipping, exponential, and the min-max transformations (the first three columns respectively), together with the firing rate (right column) for the three-cluster SBM. Here *t_R_*= 0.005 s. Other parameters are set as following Tab. S1 with *T* = 20 s, Δ*t* = 0.0001 s. Higher *p*_out_ and stronger inhibition generally reduce recovery by making the planted communities less dynamically distinct. Differences in *f*_1_ and *f*_2_ restores separation, yielding higher ARI away from *f*_1_ *f*_2_. The firing-rate plot indicates that changes in the recovery is not directly related to the changes in the average activity level.

**Figure S11:**
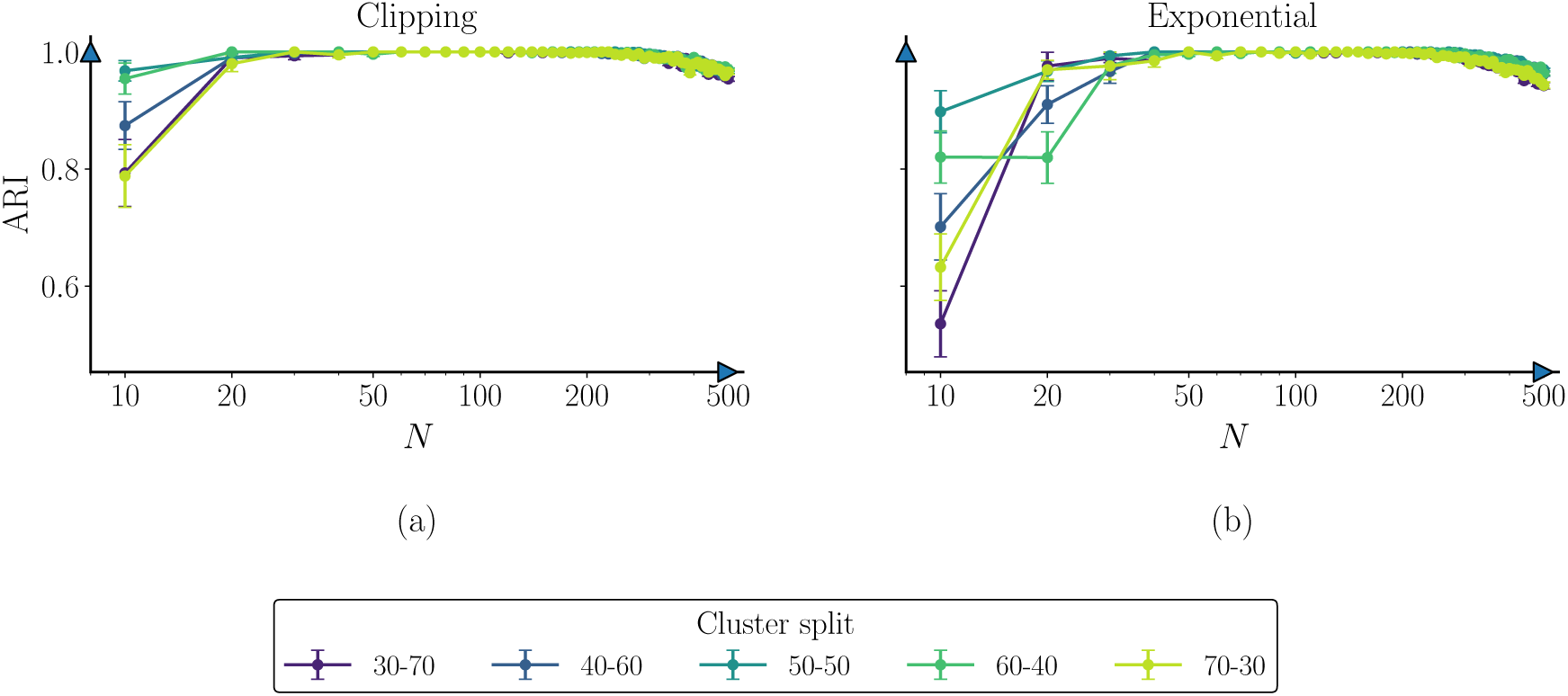
Mean ARI and firing rate vs network size *N* for the clipping transformation (a), and exponential transformation (b) for different planted two-cluster SBM splits in the conductance based LIF network model. Here *p*_in_ = 0.8, *p*_out_ = 0.01, *g_E_* = 0.5, *g_I_* = 0.1, *r_C_* = 100 Hz, and *f_C_* = 0.2, *t_R_* = 0.005 s. Both transformations achieve ARI 1 for moderate and large *N* with balanced splits slightly outperforming the imbalanced ones. Results are averaged over 20 independent simulations with standard error bars.

**Figure S12:**
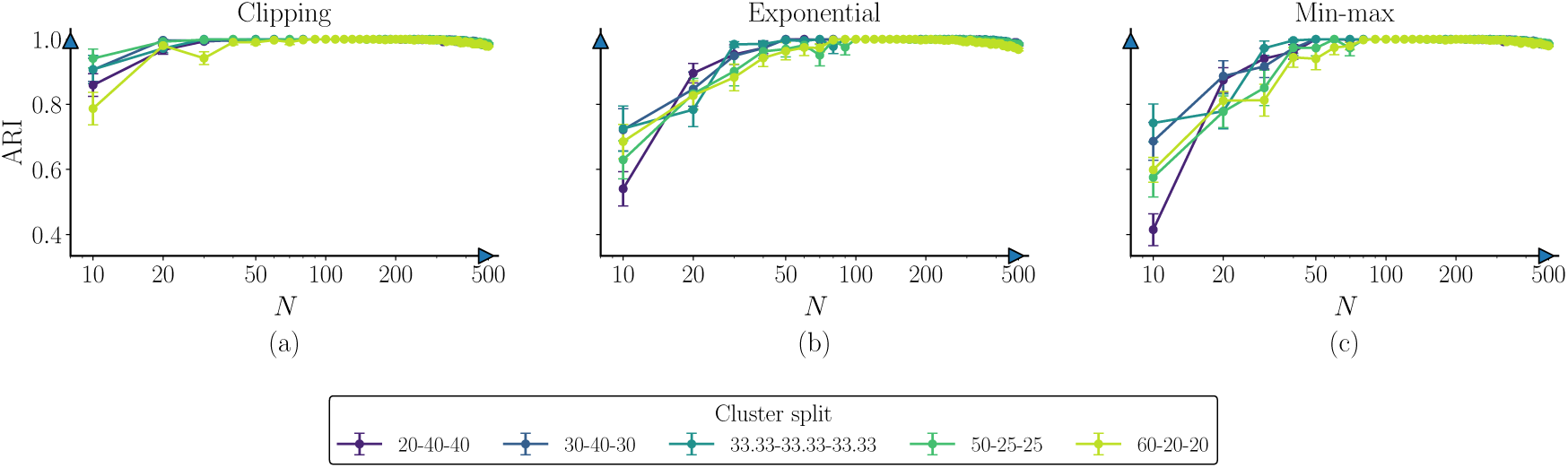
Mean ARI and firing rate vs network size *N* for the clipping transformation (a), exponential transformation (b), and min-max transformation (c) for different planted three-cluster SBM splits in the conductance based LIF network model. Here, set *p*_in_ = 0.8, *p*_out_ = 0.01, *g_E_* = 0.5, *g_I_* = 0.1, *r_C_* = 100 Hz, and *f_C_* = 0.2, *t_R_* = 0.005 s. Across all kernels ARI increases fast with *N* before saturating close to 1. More balanced cluster splits achieve higher recovery accuracy than imbalanced ones. The exponential and min-max transformations exhibit marginally higher ARI than the clipping, especially for small and intermediate *N*.

**Figure S13:**
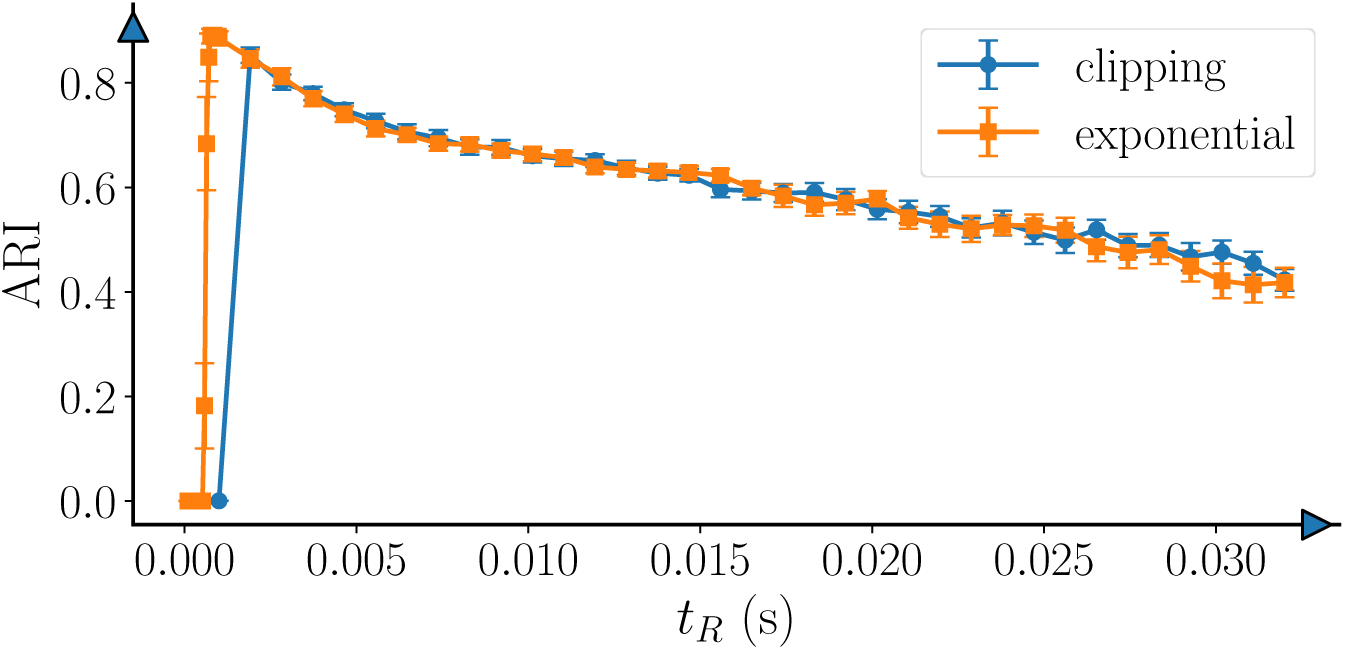
Mean ARI vs the van Rossum temporal scale *t_R_* for the conductance based LIF network. Here *t_R_* ∈ [0.0001, 0.032] s. For very small *t_R_* the ARI is poor and there exists an optimal *t_R_* ≈ 0.001 s where ARI performs the best. The ARI beyond that point decreases gradually as well

**Figure S14:**
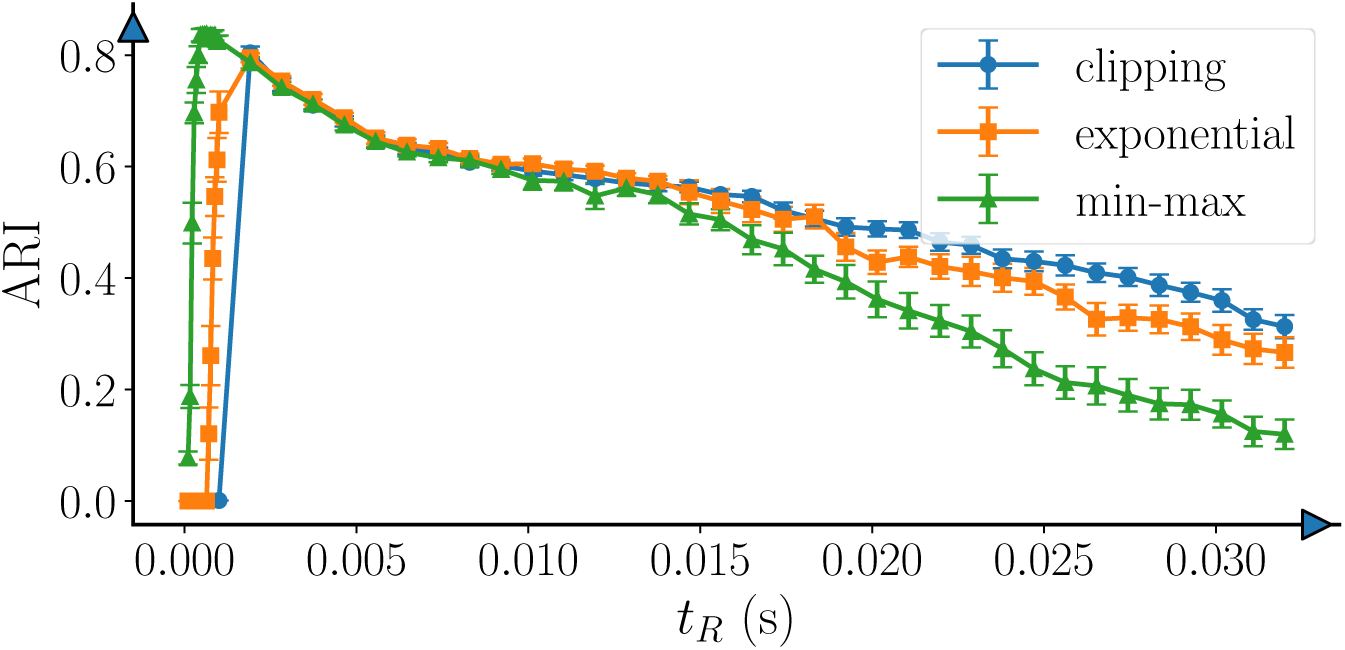
Mean ARI vs the van Rossum temporal scale *t_R_* for the conductance based LIF network. Here *t_R_* ∈ [0.0001, 0.032] s and apply our methodology. For very small *t_R_* the ARI is poor and there exists an optimal *t_R_* 0.001 s where ARI performs the best. The ARI beyond that point decreases gradually also.

**Figure S15:**
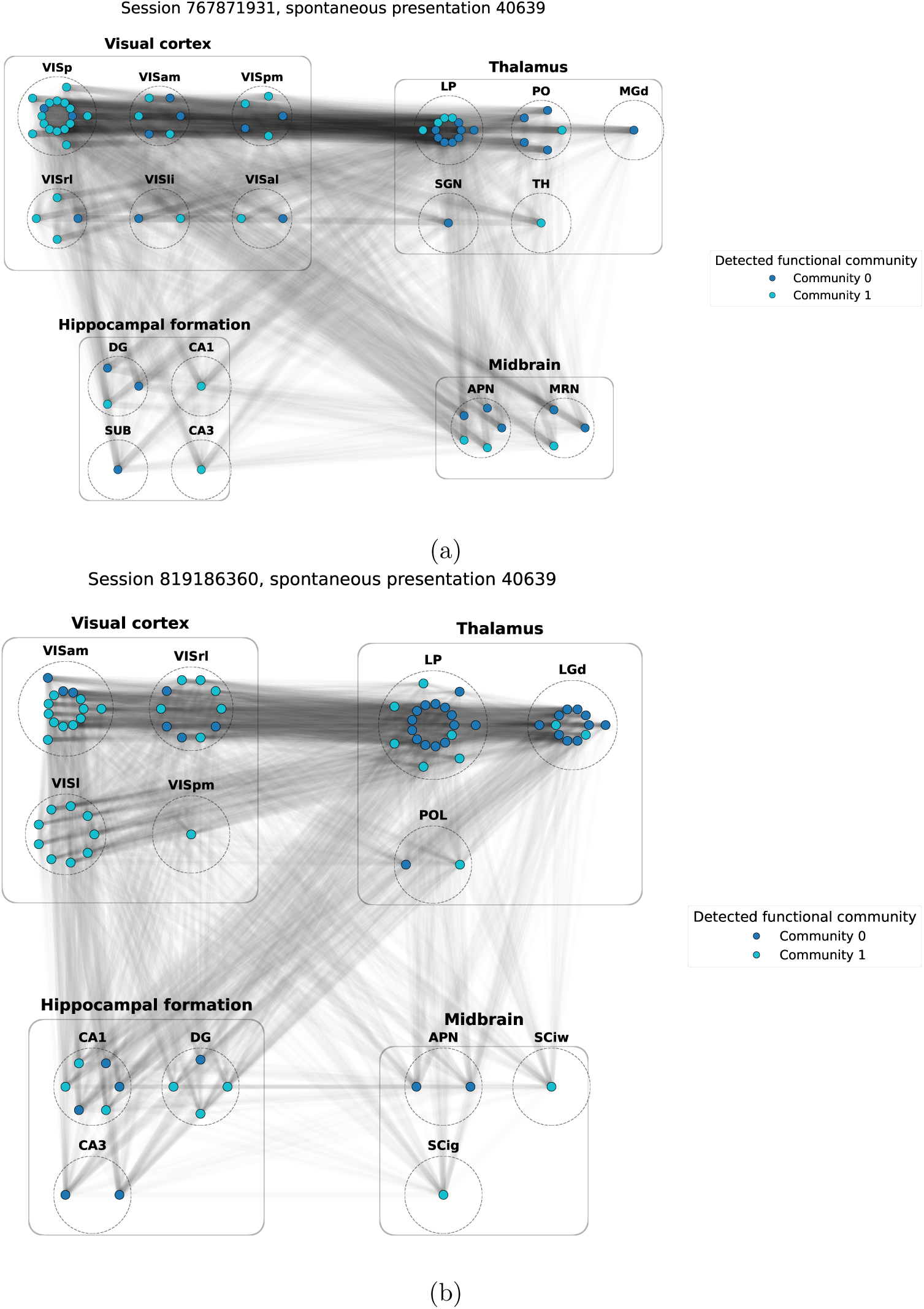
Functional neural assemblies detected using our methodology from spontaneous data in two Allen sessions: (a) 767871931 and (b) 819186360. Nodes represent neurons coloured by detected community, dashed boundaries denote anatomical brain regions including the visual cortex, thalamus, the hippocampus, midbrain, etc., and edge weights are scaled by functional similarity. The recovered communities span multiple anatomical regions while exhibiting region-specific enrichment. This suggests distributed functional organisation with partial correspondence to anatomical structure.

### Algorithm 1

Community Detection from LIF Spike Rasters Using vR-FCD

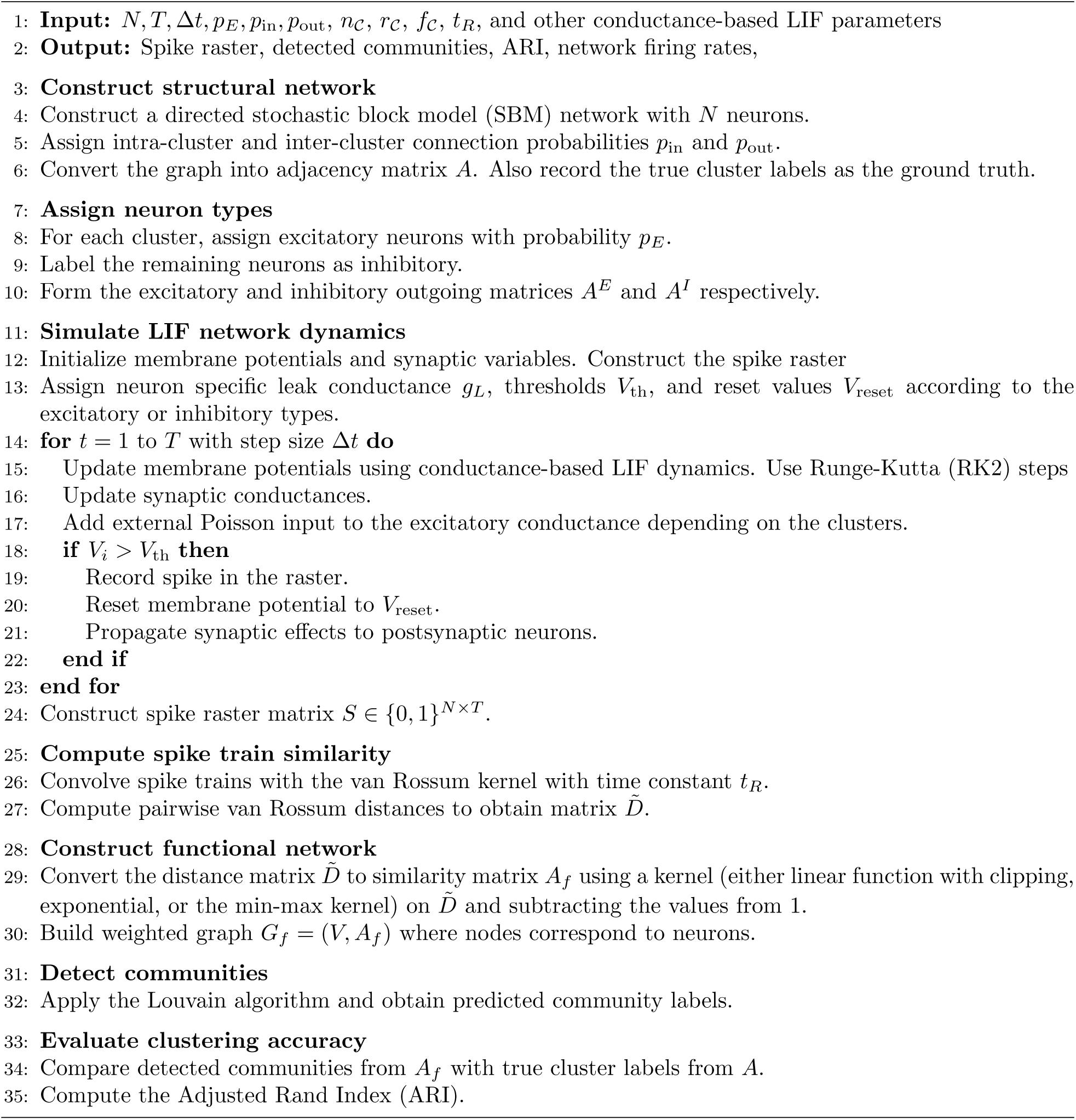

Across the two-dimensional parameter sweeps, all transformations reproduce the same qualitative dependence on the network structure and dynamics. Recovery is strongest when the planted clusters remain structurally and dynamically distinguishable, and deteriorates when stronger inter-cluster coupling or similar external inputs make their spike activity more homogeneous (Figs. S7 and S8). Repeating these sweeps under a second, more challenging parameter regime portrays the same overall picture, however with reduced recovery. Heterogenous input amplitudes can nevertheless restore separability between clusters (Figs. S9 and S10). Community recovery also improves quick with network size *N*, with balanced partitions slightly outperforming the imbalanced ones (Fig. S11 and S12). There is a slight degradation in the perfromance when *N* becomes very large from growing number of cross-community connections and stochastic interactions. Finally, all transformations show a clear dependence on the van Rossum time scale parameter *t_R_* with a poor recovery for very small value, an optimal intermediate range, and gradual deterioration for larger values (Figs. S13) and S14).

## G Application of the vR-FCD framework to the Allen Neuropixels electrophysiology data

To test the vR-FCD framework on experimental neural data (§ 5), we use recordings from the Allen institute, which has been designed to characterise large-scale neuron populations activity across cortical and subcortical regions in awake mice. The data collection used two-photon (2P) imaging to measure brain activity from labels of cortical regions in awake mice who are subjected to a range of visual stimuli. They also use Neuropixel probes to collect spike activity data from around 100,000 neurons coming from *wild-type C57BL6/J* mice and three other transgenic lines of mice: *Sst-IRES-Cre; Ai32, Pvalb-IRES-Cre; Ai32 and Vip-IRES-Cre; Ai32*. Mice were exposed to either of two possible stimulus sets: “Brain Observatory 1.1” and “Functional Connectivity”. Both consist of stimulus presentation types like “Gabors”, “Flashes”, “Drifting Gratings”, “Natural Movie One”, etc. The duration between two stimulus presentation when a mouse is not exposed to any visual stimulation is referred to as “spontaneous” activity. During this period the screen is held at a uniform mean luminance (gray), allowing measurement of the spiking activity in the absence of any sensory drive. The Brain Observatory 1.1 merges two sessions from the 2P dataset. The Functional Connectivity data consists of a subset of the Brain Observatory dataset, but with higher number of repeats. This aids researchers in studying functional interactions which is highly relevant for our work.

The Neuropixels probes consist of either 374 or 383 channels that measure the voltage fluctuations in the neurons. The data is collected as “spike-bands” (frequency of 30 kHz) and tells us about the neurons’ actions potentials which are situated adjacent to the probes. The spike-band recordings undergo further processing and are released in the *Neurodata Without Borders* format (https://nwb.org/), facilitating standardised access and reuse by the research community. These steps include subtracting the median value of the channel to center the signals around zero, removal of the median across channels to get rid of common-mode noise, high-pass filtering and whitening across blocks of 32 channels, and spike-sorting using `Kilosort2` [42] (GitHub link: https://github.com/MouseLand/Kilosort) to detect spikes from individual “units”. For the data processing steps by Allen institute. Although the processing steps include additional stages, our analysis focuses on the spike-sorted raster data. Because spike sorting does not guarantee a one-to-one correspondence between an identified cluster of spikes and a single biological neuron, the AllenSDK refers to these recorded entities as “units”, and we adopt the same terminology throughout. The spikes have some probability of getting mixed together if there are considerable movement of the neurons compared to the probes or a neuron is far away from the probe.

In this paper we restrict our analysis to “spontaneous” stimulus presentations, when the subjects are not exposed to any visual stimuli, letting us work with internally generated activity patterns in the neural population rather than the stimuli driven ones. This also helps avoid confounding spike-train similarity due to the stimulus responses, because visual stimuli can induce transient correlations across neurons, which we do not want for our analysis. Spontaneous activity, on the other hand, is more robust in terms of the intrinsic population organisation. We further restrict our analysis to the wild type C57BL6/J mice. The reason is three-fold:

i. the majority of the recordings are from the wild type C57BL6/J mice, providing the largest and most consistent subset for cross-session comparisons,
ii. provides a more homogeneous baseline populations and avoids confounding effects associated with cell-type specific targeting; and
iii. excluding transgenic lines reduces additional biological and experimental heterogeneity that could influence comparisons of functional community structure across sessions.

Recordings from the wild type mice samples provide a consistent baseline for comparisons across sessions. We therefore focus on the Functional Connectivity stimulus set, which contains longer repeated spontaneous recordings and sufficient spike activity for reliable similarity estimation and community detection. The retained sessions and spontaneous presentations are summarised in tables S2 and S3, which report the session-level metadata, recording scale, and the durations of the spontaneous activity segments used in the study.

**Table S2:** Allen SDK extracellular electrophysiology (ecephys) data.

| Session ID | Specimen ID | Age (days) | Sex | Unit count | Channel count | Probe count |
| --- | --- | --- | --- | --- | --- | --- |
| 766640955 | 744912849 | 133.0 | M | 842 | 2233 | 6 |
| 767871931 | 753795610 | 135.0 | M | 713 | 2231 | 6 |
| 768515987 | 754477358 | 136.0 | M | 802 | 2217 | 6 |
| 771160300 | 754488979 | 142.0 | M | 930 | 2230 | 6 |
| 771990200 | 756578435 | 108.0 | M | 546 | 2229 | 6 |
| 774875821 | 759711152 | 114.0 | M | 649 | 2233 | 6 |
| 778240327 | 760938797 | 120.0 | M | 784 | 2234 | 6 |
| 778998620 | 759674770 | 121.0 | M | 793 | 2229 | 6 |
| 779839471 | 760960653 | 122.0 | M | 863 | 2220 | 6 |
| 781842082 | 760946813 | 126.0 | M | 833 | 2232 | 6 |
| 793224716 | 769319624 | 120.0 | M | 781 | 2229 | 6 |
| 819186360 | 800249587 | 128.0 | F | 531 | 1696 | 5 |
| 821695405 | 800250057 | 134.0 | F | 474 | 1856 | 5 |
| 847657808 | 827809884 | 126.0 | F | 874 | 2298 | 6 |

**Table S3:**
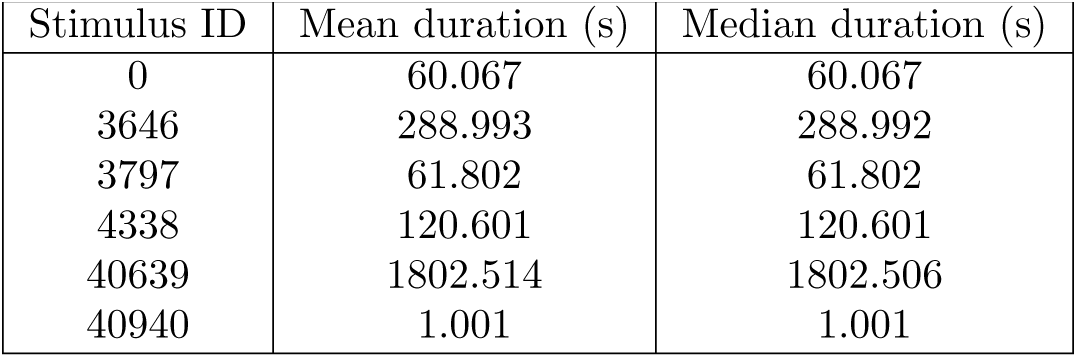
Stimulus IDs with mean and median durations in seconds.

To study the anatomical correspondence of the detected functional assemblies, we compare the communities detected from stimulus ID 40639 using the vR-FCD framework with the recorded brain region labels. The labels used for region identification followed from Siegle *et al.* [28] are as follows: “visual cortex (VISp, VISl, VISrl, VISam, VISpm, VISal, VISmma, VISmmp, VISli, VIS), thalamus (LGd, LD, LP, VPM, TH, MGm, MGv, MGd, PO, LGv, VL, VPL, POL, Eth, PoT, PP, PIL, IntG, IGL, SGN, VPL, PF, RT), hippocampal formation (CA1, CA2, CA3, DG, SUB, POST, PRE, ProS, HPF), midbrain (MB, SCig, SCiw, SCsg, SCzo, SCop, PPT, APN, NOT, MRN, OP, LT, RPF), and other/nonregistered (CP, ZI, grey)”. Representative results are shown for session IDs 767871931 and 819186360.

